# Increased prevalence of a frontoparietal brain state is associated with better motor recovery after stroke affecting dominant-hand corticospinal tract

**DOI:** 10.1101/2022.02.10.479962

**Authors:** Emily Olafson, Georgia Russello, Keith W Jamison, Hesheng Liu, Danhong Wang, Joel E Bruss, Aaron D Boes, Amy Kuceyeski

## Abstract

Strokes cause lesions that damage brain tissue, disrupt normal brain activity patterns and can lead to impairments in motor function. Although modulation of cortical activity is central to stimulation-based rehabilitative therapies, aberrant and adaptive patterns of brain activity after stroke have not yet been fully characterized. Here, we apply a brain dynamics analysis approach to study longitudinal brain activity patterns in individuals with ischemic pontine stroke. We first found 4 commonly occurring brain states largely characterized by high amplitude activations in the visual, frontoparietal, default mode, and motor networks. Stroke subjects spent less time in the frontoparietal state compared to controls. For individuals with dominant-hand CST damage, more time spent in the frontoparietal state from 1 week to 3-6 months post-stroke was associated with better motor recovery over the same time period, an association which was independent of baseline impairment. Furthermore, the amount of time spent in brain states was linked empirically to functional connectivity. This work suggests that when the dominant-hand CST is compromised in stroke, resting state configurations may include increased activation of the frontoparietal network, which may facilitate compensatory neural pathways that support recovery of motor function when traditional motor circuits of the dominant-hemisphere are compromised.

## 1 Introduction

The ability to perform motor functions after stroke depends on the coordinated reconfiguration of distinct global brain activity patterns (Park et al., 2011). Novel data-driven methods to characterize brain activity at single TR resolutions have illuminated the dynamic nature of brain activity in several brain disorders (Yang et al., 2021; Braun et al., 2021; Adhikari et al., 2020; Kaiser et al., 2019; Marshall et al., 2020; Parker Singleton et al., 2021), but these approaches have not yet been applied to stroke patients and related to functional recovery. These methods provide information about the dynamics of brain function complementary to and beyond traditional static measures of functional connectivity (Cornblath et al., 2020), and thus may provide new insights regarding the process of recovery following stroke.

Prior work characterizing spatiotemporal brain dynamics after stroke has focused on identifying altered functional connectivity states, which reflect time-varying patterns of functional connectivity (FC). So-called “dynamic FC” analyses identify recurrent connectivity patterns using a sliding window approach, in which FC is repeatedly calculated over consecutive windowed segments of the fMRI scan. This approach yields FC networks that fluctuate over time, with a temporal resolution proportional to the size of the window; about 30-60 seconds (Savva et al., 2019). In stroke populations, dynamic FC studies have demonstrated stroke-related differences in temporal configurations of motor networks (Bonkhoff et al., 2020a) and participation in connectivity states that varies with severity (Bonkhoff et al., 2020b).

In contrast, single-TR analysis of activation states identified in a data-driven fashion using k-means clustering of the time series data (Cornblath et al., 2020) provides a closer look at the moment-to-moment changes in recurrent brain activity, with the time spent in each state lasting, on average, 5-10 seconds (Cornblath et al., 2020). A benefit to analysing brain activation states over or alongside connectivity states is that activation patterns can enable a more refined interpretation of connectivity differences between groups. FC is traditionally defined as the correlation between two brain region’s activity over time. FC may be driven by two distinct features of brain activity: by the individualized spatial patterns of large-amplitude activations (Zamani Esfahlani et al., 2020), and by the amount of time spent in recurring patterns of activity (Betzel et al., 2021; Baker et al., 2014). In this paper we aim to identify group-level patterns of brain activity after stroke that relate to recovery, and assume that recurring activity patterns are shared across individuals but are expressed in different proportions. Understanding the temporal patterns of activity underlying recovery-relevant FC changes after stroke can aid in the development of more accurate targets in stimulation therapies. To our knowledge, this fine-grained view of activation state changes after stroke has not yet been explored.

In this paper, we aim to identify patterns of activation that are associated with better motor recovery. Recent work has highlighted the importance of frontoparietal areas in supporting motor abilities in the chronic phase of stroke (Tscherpel et al., 2020) specifically in patients with poor corticospinal tract (CST) integrity (Hordacre et al., 2021a). When these descending motor pathways are significantly damaged, descending white matter tracts from higher-order motor areas, such as the premotor cortex, may support motor output. Greater white matter tract integrity is found in frontoparietal tracts of the dominant hemisphere (i.e., the hemisphere corresponding to the dominant limb) in healthy individuals (Howells et al., 2018), attributable to programs and skills built over long periods of dominant limb use. Individuals with dominant hemisphere CST damage may therefore more easily recruit frontoparietal networks to support motor recovery compared to individuals with non-dominant hemisphere CST damage. Understanding this type of subject- and lesion-specific post-stroke recovery process is precisely the type of information needed to develop personalized rehabilitation strategies to maximally promote recovery.

Here, we propose to first identify and characterize recurring brain activity patterns, or states, in healthy controls and individuals with ischemic pontine stroke (Cornblath et al., 2020). We hypothesized that individuals with ischemic stroke would display altered dynamic brain state metrics, e.g. fractional occupancy, dwell time and appearance rates, compared to control subjects, and, further, that these dynamic state metrics would be associated with measures of motor recovery. We specifically hypothesized that increased occupancy in a brain state characterized by frontoparietal activation would be related to better later-stage motor recovery in stroke subjects with damage to their CST in the dominant-hand hemisphere. Finally, to bridge dynamic brain state analyses and more classic functional connectivity approaches, we assessed the relationship between the amount of time spent in different brain states and the FC between several resting-state networks. This last analysis is particularly important in terms of linking our current findings to previous studies of how rehabilitation techniques, including non-invasive brain stimulation, modulate the functional connectome and possibly motor recovery.

## 2 Materials and methods

### 2.1 Data description

The data consist of 23 first-episode stroke patients (34-74 years old; mean age 57 years; 8 female) with isolated pontine infarcts and 24 healthy age- and sex-matched controls (33-65 years old; mean age 52 years; 10 female). A subset of the data (11 stroke subjects and 11 healthy control subjects) used here has been previously described in Lu et al., 2011; the current study includes an additional 12 stroke subjects and 13 control subjects. Of the twenty-three stroke subjects, fourteen had right brainstem infarcts and nine had left brainstem infarcts (Figure S1 and S2). Patients were scanned between two and five times over a period of 6 months. Specifically, MRIs were obtained at 7, 14, 30, 90 and 180 days after stroke onset on a 3T TimTrio Siemens using a 12-channel phase-array head coil. Fugl-Meyer assessments were performed twice for each subject at each session and averaged. The Fugl-Meyer test includes 33 tasks that assesses motor function, balance, sensation, and joint function of the upper limbs (Fugl-Meyer et al., 1975). Each task was rated on a scale of 0 to 2 (0 indicates the subject was unable to perform the task, 1 indicates the subject could partially perform the task, and 2 indicates the subject was able to perform the task). The total sum of the 33 scores was then normalized to a score between 0 and 100, where 100 represents the best possible performance across all 33 tasks. Anatomical images were acquired using a sagittal MP-RAGE three-dimensional T1-weighed sequence (TR, 1600ms; TE 2.15ms; flip angle, 9°, 1.0 mm isotropic voxels, FOV 256 × 256). Each MRI session involved between two and four runs of task-free fMRI at 6 minutes each. Subjects were instructed to stay awake with their eyes open; no other task instruction was provided. Images were acquired using the gradient-echo echo-planar pulse sequence (TR, 3000ms; TE, 30ms; flip angle, 90°, 3 mm isotropic voxels). Anatomical MRI, lesion masks and fMRI data were processed as described below and in (Olafson et al., 2021).

### 2.2 Anatomical MRI processing

Preprocessing of the longitudinal anatomical MRIs included affine registration of each subject’s T1 scans to the baseline T1 scan, collapsing co-registered files to an average T1 and creation of a skull-stripped brain mask followed by manual editing and binarization of the hand-edited mask. The brain mask was then transformed back to each of the follow-up T1s in native space using the inverse registration acquired from the first step. This was followed by bias field correction of all the T1 scans, transformation of native-space bias field-corrected data back to baseline space, and the creation of an average bias field-corrected scan for each subject. Stroke lesion masks were hand-drawn on these transformed T1 scans by ADB and JEB. Structural normalization was performed with the ANTs toolbox (Avants et al., 2011).

### 2.3 Functional MRI processing

Preprocessing of the longitudinal functional MRIs was performed using the CONN toolbox (Whitfield-Gabrieli and Nieto-Castanon, 2012), including functional realignment of volumes to the baseline volume, slice timing correction for alternating acquisition, segmentation and normalization, and smoothing with a 4 mm FWHM kernel. This was followed by a denoising protocol (CompCor) (Behzadi et al., 2007) which regressed out the cerebrospinal fluid and white matter signal, as well as 24 realignment parameters (added first-order derivatives and quadratic effects). Temporal band-pass filtering (0.008 - 0.09 Hz), despiking and global signal removal regression were also performed. The first four frames of each BOLD run were removed. Frame censoring was applied to scans with a framewise displacement threshold of 0.5 mm along with its preceding scan (Power et al., 2012). Regional time series were acquired by parcellating the scans into 268 non-overlapping brain regions using a functional atlas derived from healthy controls (Shen et al., 2013) and averaging the time course of all voxels within a given region. Voxels identified as lesioned were excluded from regional time series calculations. The first 200 volumes from each subject’s fMRI were used for subsequent analyses to ensure equal contribution of each scan to the brain state clustering (see below for details). Finally, each of 268 regions was assigned to one of 8 functional networks, identified by (Finn et al., 2015) using spectral clustering in healthy subjects (Figure S3) named as follows: medial frontal network, frontoparietal network, default mode network, subcortical/cerebellum network, motor network, visual I network, visual II network, and the visual association network. These networks reflect collections of brain regions whose temporal signals are homogeneous at rest (i.e., the activity of regions within each network is similar over time) in a healthy population, and are referred to as “canonical” networks due to their repeated observation in resting-state data.

### 2.4 Dynamic brain states and their metrics

Following Cornblath et al. (2020), all subjects’ regional fMRI time series were concatenated, producing an n x p matrix where n = 47 subjects * 200 TRs * 2-5 sessions and p = 268 brain regions). This matrix was z-scored along columns such that each brain region had a mean of 0 and a standard deviation of 1. K-means clustering was then applied to identify clusters of brain activation patterns, or states (Figure 1A). Pearson correlation was used as the cluster distance metric and clustering was repeated 50 times with different random initializations before choosing the solution with the best separation of the data (minimum sum of point-to-centroid distances, summed over all k clusters). To determine the optimal number of clusters and evaluate the quality of clustering, we performed several analyses (Figure S4). First, we plotted the variance explained by clustering (between-cluster variance divided by the sum of between-cluster and within-cluster variance) for k = 2 to 12 and identified the curve’s “elbow” at k = 4 as a potential optimal number of clusters. We also plotted the distortion curve, which is the average distance from each point to its centroid and again determined via elbow criteria an optimal cluster number of 4. We then plotted silhouette coefficients for k = 4 to assess if there was evidence of misassignment of any of the points. To further assess the stability of clustering and ensure our partitions were reliable at k = 4, we repeated the above clustering process 50 times and compared the adjusted mutual information (AMI) between each of the 50 results. The partition which shared the greatest total AMI with all other partitions was selected as the final cluster assignment. The centroids of each state (cluster) were calculated by taking the mean of all TRs assigned to that state in regional activation space (Cornblath et al., 2020). Following Cornblath et al., dominant networks in each state were determined by calculating the cosine similarity between each of 8 networks and each centroid. High and low amplitude network-level activations were assessed separately by taking the cosine similarity of the positive and negative parts of the centroid (and zeroing out values with the opposite sign), respectively.

**Figure 1:**
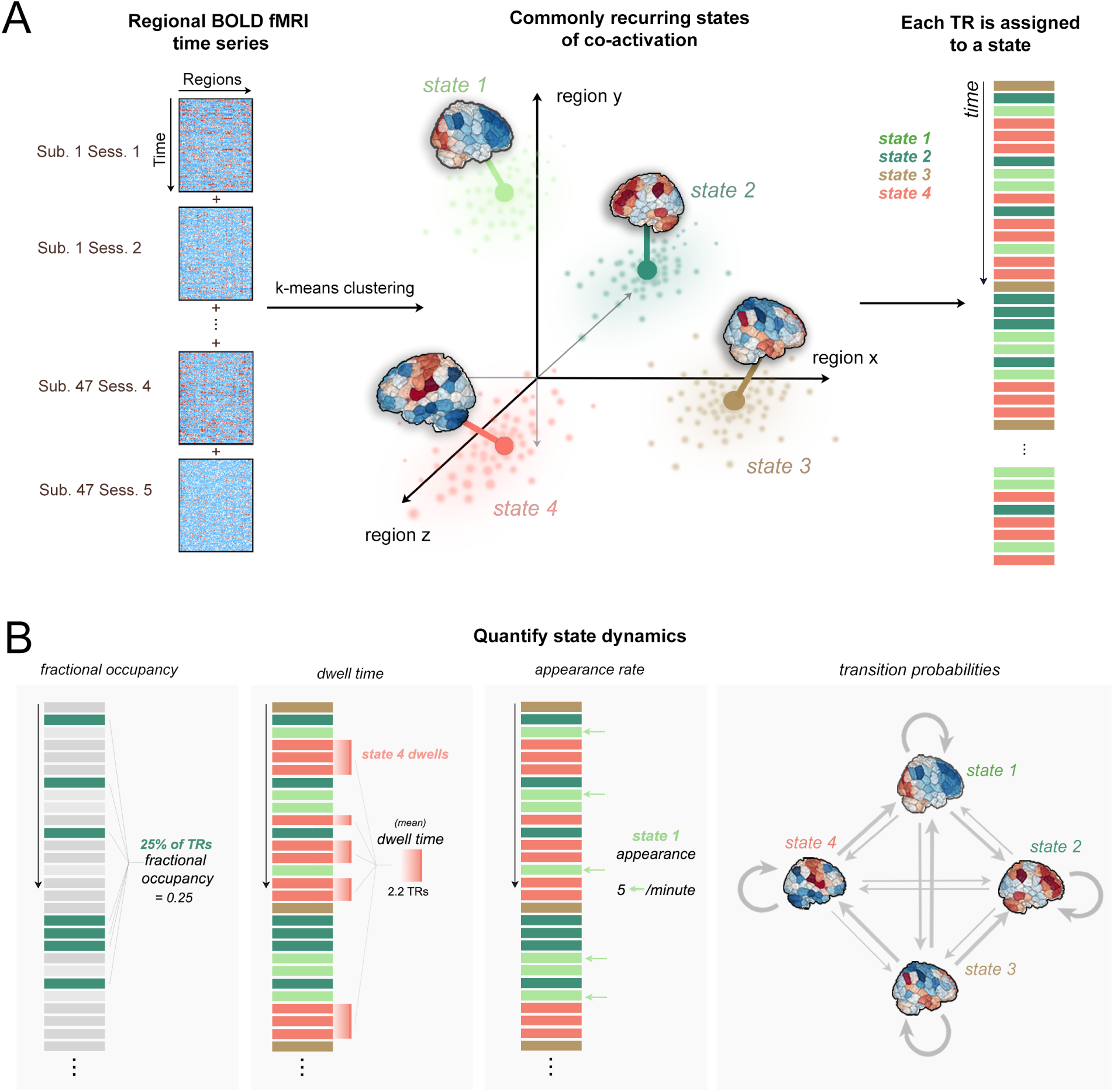
Clustering of time series data and quantification of dynamic state metrics. **A**. Time series data from all subjects were concatenated together along the time dimension. K-means clustering produced 4 distinct brain activation states defined by different locations in regional activation space (image adapted from Cornblath et al., 2020). Each TR is assigned to one of four brain states based on k-means partitions. **B**. Fractional occupancy, dwell time, appearance rate, and transition probabilities are calculated separately for each subject and for each state.

We performed several analyses to assess the robustness of our results under different conditions. First, we repeated the entire clustering process using two other brain atlases of varying resolutions: the group average FreeSurfer Desikan-Killany atlas with additional cerebellum and subcortical regions (86 regions) (Desikan et al., 2006) and the CC400 atlas (400 regions) (Craddock et al., 2012). Finally, we performed the clustering after combining stroke and control data together because we were interested in determining differences in shared activation states across both groups. However, it may be possible that stroke and control subjects occupy distinct states that can only be observed by clustering stroke and control subjects separately. Therefore, we repeated the clustering on the stroke and control subject data separately to determine if there were any differences in the resulting states compared to those obtained with the combined data.

Four brain states were identified and characterized based on the activity of canonical resting-state networks with the states (see Results): one state with high frontoparietal (FPN) activation, one with low frontoparietal (FPN) activation, one with high motor network activation, and one with low motor network activation. Once the brain states were determined, several metrics were calculated to characterize each individuals’ dynamics (Figure 1B):

1. **Fractional occupancy (FO):** probability of occurrence of each state (TRs in a state/total TRs).
2. **Dwell time (DT):** average time spent in each state before switching states (avg. # consecutive TRs assigned to each state)
3. **Appearance rate (AR):** number of times a state it transitioned into per minute.
4. **Transition probabilities (TPs):** probability of switching to any given state i from state j (includes probability of transitioning to the same state)

FO, DT, AR and TPs were calculated separately for each of the 5 sessions (1 week, 2 weeks, 1 month, 3 months, and 6 months post baseline or stroke) and the average FO/DT/AR/TPs for each subject was obtained by taking the mean across their longitudinal sessions. State dynamics metrics were compared between groups using unpaired t-tests and corrected for multiple comparisons using Benjamini-Hochberg (BH) and false discovery rate (FDR) of 0.05 (Benjamini and Hochberg, 1995).

### 2.5 Assessment of corticospinal tract integrity

The probabilistic Tang brainstem atlas (Tang et al., 2018) was used to define left and right binary CST masks in 2mm MNI space by voxel-wise thresholding at 50%. The Dice overlap between the left and right CST masks and the binarized lesion mask was calculated for each subject and lesions were visualized to verify intersection with the CST (Figure S1 and S2). In one subject (SUB13), Dice overlap with CSTs were low; however, upon visualization the lesion appeared to impact CST ventral to the atlas, i.e. in the spinal cord. This was confirmed by assessing the lesion’s overlap with a spinal cord atlas (Figure S10) using the Spinal Cord Toolbox (De Leener et al., 2017). In total, 21/23 subjects had CST damage and 9/23 subjects had damage to their dominant-hand CST, but only 8/23 had motor assessment data at the 3- and 6-month time points. Of the 9 subjects with dominant CST damage, 8/23 were right-handed and either had left CST damage superior to decussation or right spinal cord damage inferior to decussation, and 1/23 was left-handed and had right CST damage.

### 2.6 Relating *FPN*^+^ state metrics to 3- and 6-month motor performance outcomes

We were interested in determining whether *FPN*^+^ state metrics related to longer-term (3-6 months post-stroke) motor performance outcomes specifically in subjects with dominant CST damage. To assess the interaction effect of hand dominance on the relationship between the frontoparietal state parameters (FO and DT) over time and Fugl-Meyer (FM) scores over time, four linear models were constructed:

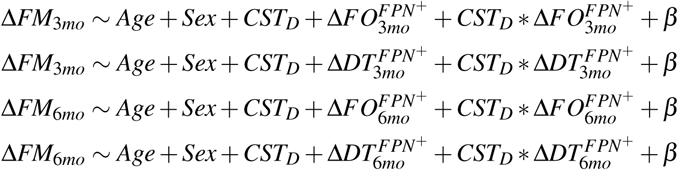

Where:

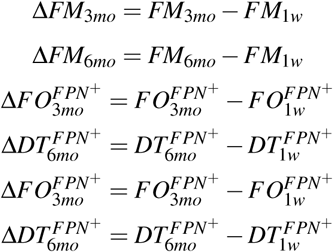

and *CST*_*D*_ is a binary variable indicating whether the subject’s dominant-hand CST had non-zero Dice overlap with the lesion. The models were constructed using only the FPN+ metrics that were found to have significant differences between stroke and controls at the 3 and 6 month time points. P-values obtained for each predictor across all models were corrected for multiple comparisons using BH-FDR and a threshold of 0.05.

### 2.7 Effect of proportional recovery on observed relationships

We were interested in determining whether relationships observed in the above model were driven by the correlation (if any) between DT/FO in *FPN*^+^ and baseline FM scores (1-week post-stroke) in subjects with dominant-hand CST damage. Because most subjects obtained proportional recovery or greater, i.e. their final (6 month) impairment was *≥* 70% of their initial impairment (Kundert et al., 2019), their 1-week FM scores were strongly correlated with 3- and 6-month FM scores (a phenomenon called the ceiling effect, see Hope et al., 2019). Therefore, it is possible that relationships between Δ*FM* and 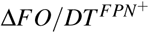 observed in the linear model could be driven by the underlying relationship between baseline FM scores and 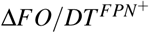. To explore this possibility, for those *FPN*^+^ metrics whose change had a significant correlation with baseline FM (which were 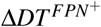 and 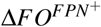 for 3 and 6 months), we performed permutation testing to obtain the null distribution of correlations between Δ*FM* and 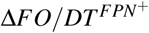, assuming patients obtain only proportional recovery (PR). *FM*_*initial*_, 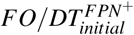 and 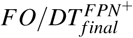 were fixed to their actual observed values, but 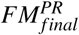 was randomly generated via a parametric bootstrap by simulating scores according to the proportional recovery rule as in Kundert et al., 2019. Specifically, each subject’s 3- and 6-month FM scores 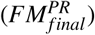 were set to 70% of their initial impairment (100-*FM*_*initial*_) with an noise term *ε* ∼ *N*(0, 3):

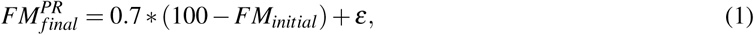

The p-value for the correlation between the observed Δ*FM* and 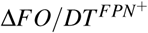 and was then calculated by calculating the proportion of times the null correlation exceeded the true correlation (Figure S11C,D). If that p-value is significant, then we can be more confident that 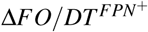 is correlated with the change in FM above and beyond baseline FM and the expected by proportional recovery.

### 2.8 Comparison of fractional occupancy and functional connectivity

Finally, we wanted to understand how differences in state dynamics between controls and stroke subjects could translate to differences in FC between the two groups. We hypothesize that individuals with more TRs (higher FO) in a brain state with high co-activation of two networks will result in larger positive FC between those networks, while more TRs (higher FO) in a brain state with large activations in opposite directions (contra-activation) of two networks will result in a more negative FC between those networks (Figure S12). We tested this hypothesis for our brain state of interest, *FPN*^+^, in the following way. First, we identified the pairs of networks that were highly co-activated/contra-activated during *FPN*^+^, which was defined as having an absolute value cosine similarity with the centroid of *FPN*^+^ of greater than 0.2 (chosen heuristically as the threshold separating networks active vs. not active during a given state). We only analyzed the networks with larger magnitude co-activations/contra-activations since the networks with activity closer to zero in the *FPN*^+^ state are not likely to be influenced by changes in FO of this state. We first calculated the functional connectivity as the Pearson’s correlation between each pair of 268 regions and performed a Fisher’s r-to-z transformation of the FC weights. We then averaged the FC values between regions belonging to each pair of networks to produce a network-level FC (i.e., the average FC within each of 8 predefined networks, including the frontoparietal network). We correlated each subject’s FO in the *FPN*^+^ state with the FC between each pair of networks determined to be highly co-activated/contra-activated in the *FPN*^+^ state. We further demonstrated that temporal fluctuations in 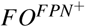 over segments of the scan are related to dynamic FC in areas belonging to the frontoparietal network. We first identified regions with high z-score BOLD signal in the *FPN*^+^ centroid (greater than 0.4), and, for those regions, calculated their sliding-window dynamic FC and 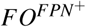 using the same window and overlap (window size = 45 seconds, overlap = 3 seconds). We then correlated these two values over the entire fMRI scan for each individual. To ensure any observed relationship was not driven by global BOLD fluctuations, we recalculated this correlation using 100 randomly selected regions’ dynamic FC as a null comparison.

#### Code availability

The code to replicate this analysis is available on GitHub: https://github.com/emilyolafson/dynamic-brainstates

## 3 Results

### 3.1 Clustering reveals an optimum of 4 brain states

We observed the change in variance explained with increasing k (Figure S4A); the difference in variance explained between k = 4 and k = 5 was < 2 %. The elbow of the curves were between 4-6 clusters across the cluster quality metrics (Figure S4C). We chose k = 4 for parsimony, and replicate k = 5 in the supplemental information section (Figure S9. Silhouette values for k = 4 revealed good cluster assignment with very few TRs assigned erroneously (i.e. having negative Silhouette values) (Figure S4B). In general, we found that the mutual information shared between partitions was quite high (0.88-1), suggesting consistent clustering across independent initializations of k-means (Figure S4D). We replicated the Shen atlas’s 4 activity states with 86 and 400 region atlases (Figure S6). Finally, the states identified by clustering stroke and control groups separately are similar to the states identified by clustering all individuals together (Figure S5). Therefore, we report the full clustering results in the main text and include the various replications in the Supplemental Information.

### 3.2 Brain states have distinct patterns of canonical resting-state network activation

The four brain states consist of distinct activation levels across the various canonical resting-state networks (Figure 2A, B). State 1 (*FPN*^+^) was characterized by high amplitude activation of regions in the frontoparietal network and low amplitude activations in the visual I network, state 2 (*FPN*^*−*^) by low amplitude activation in the frontoparietal network and high amplitude in the visual I network (equal magnitude activations), state 3 (*MOTOR*^+^) by high amplitude activation of the somatomotor network and low amplitude activation in the default mode network, while state 4 (*MOTOR*^*−*^) by low amplitude activation of the motor network as well as high amplitude activation of the default mode network. The correlations between the 268-region centroids of the *FPN*^+^ and *FPN*^*−*^ states and the *MOTOR*^+^ and *MOTOR*^*−*^ states were strong and negative, suggesting that brain states are likely hierarchically organized into 2 meta-states that are each composed of two sub-states containing opposing activation patterns (Figure 2C), a finding which has been previously reported in other work using this technique (Parker Singleton et al., 2021). The same dominant networks are observed for states derived using atlases of varying resolution (FreeSurfer 86 region atlas, Shen 268 region atlas, and CC400) (Figure S6), though the patterns of overlap for all canonical networks are slightly different due to the different networks used to annotate those atlases.

**Figure 2:**
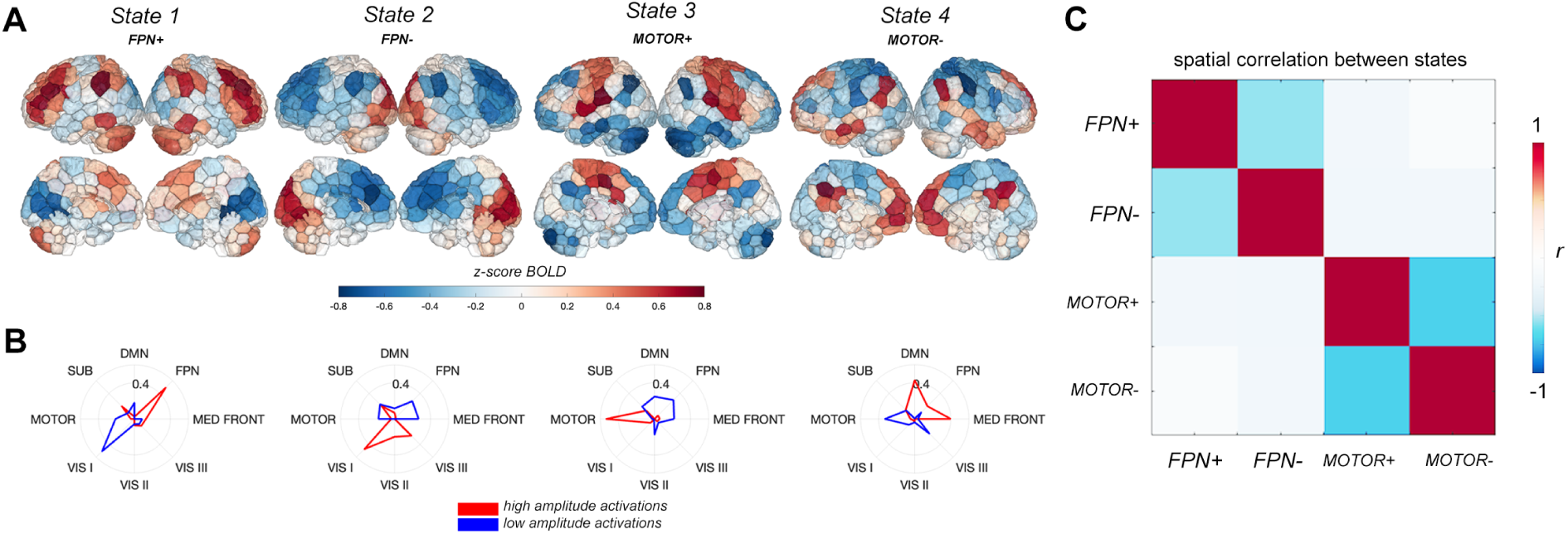
Four brain activity states. **A**. Centroids of each state calculated as the mean of the normalized regional activation over all TRs assigned to that state. **B**. Radial plots displaying cosine similarity between each cluster each canonical network; red indicates cosine similarity of the high amplitude activations, blue indicates cosine similarity of the low amplitude activations. Labels for each state were derived from analyzing the magnitude and type of overlap of each centroid with the 8 canonical networks. **C**. Region-level correlation of each pair of brain state centroids. DMN = default mode network, FPN = frontoparietal network, MED FRONT = medial frontal network, VIS III = visual association network, VIS II = visual network 2, VIS I = visual network 1, MOTOR = motor network, SUB = subcortical/cerebellum network.

### 3.3 Group differences in brain state dynamics

Fractional occupancy (FO), dwell time (DT), and appearance rate (AR) of each state were calculated for each subject. The average FO, DT, and AR across all available time points (1 week, 2 weeks, 1 month, 3 months and 6 months) for each of the four states were compared between stroke and control subjects via unpaired t-tests (Figure 3, see Supplementary Figure S7 for full session-specific results for each state, and see Supplementary Figure S8) for the general pattern of brain dynamics observed in stroke and control subjects). Stroke subjects had significantly lower FO in *FPN*^+^ compared to control subjects (p(FDR) = 0.0021), which was possibly driven more by the significantly lower dwell times in *FPN*^+^ observed in stroke subjects (p(FDR) = 0.0105) (Figure 3B, D). The frontoparietal network used in this atlas contains nodes in the dorsolateral prefrontal cortex, posterior parietal cortex, as well as nodes in the posterior inferior temporal lobe and the inferior cerebellum. Significantly smaller FO in *FPN*^+^ in stroke subjects was also observed at every session from 1 week to 6 months post-stroke (Figure 3A, with differences in *FPN*^+^ DT observed only in the chronic stage of stroke (6 months post-stroke) (Figure 3C). No differences in appearance rate of *FPN*^+^ were observed over the sessions (Figure 3E, F).

**Figure 3:**
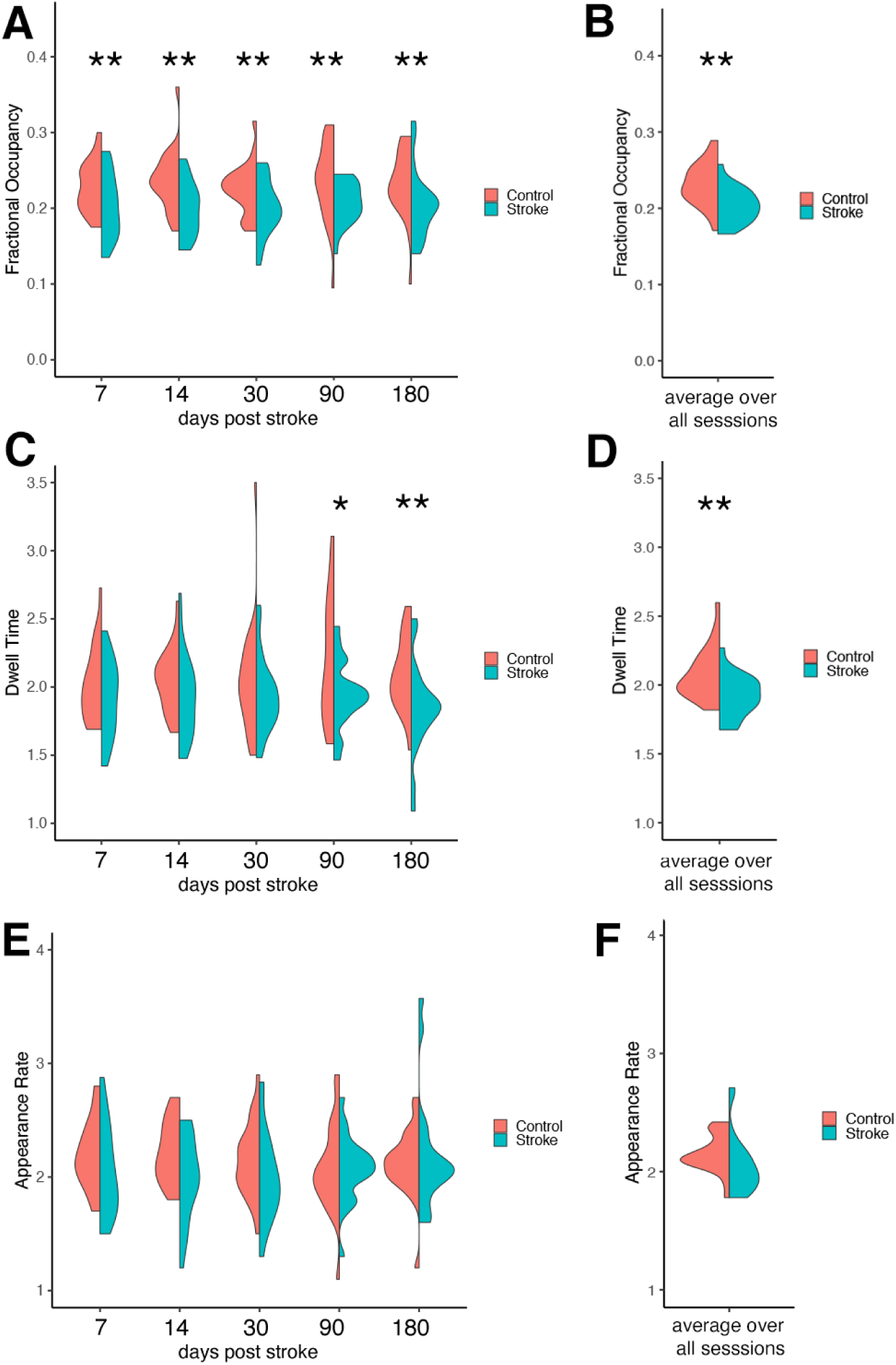
Group differences in brain state dynamics for *FPN*^+^. **A, C, E**. Stroke-control differences in session-specific fractional occupancy (FO), dwell time (DT), and appearance rate (AR) in *FPN*^+^ over the post-stroke recovery period. **B, D, F** Stroke-control differences in average FO, DT, and AR (averaged over each subject’s 2-5 longitudinal sessions). Double asterisks represent p-values < 0.05 after multiple comparisons correction, single asterisks represent significant uncorrected p-values < 0.05.

Stroke subjects had significantly reduced transition probabilities from *FPN*^+^ and *MOTOR*^*−*^ into *FPN*^+^ (p(FDR) = 0.023 and 0.020, respectively) (Figure 4A, B). The lower persistence probability in stroke for *FPN*^+^ seems to be driven by chronic-stage differences; in the session-specific analysis, individuals with stroke have significantly lower persistence probability for *FPN*^+^ at the 6-month time point (p(FDR) = 0.030) and a trend for lower persistence probability at 3 months (Figure 4C).

**Figure 4:**
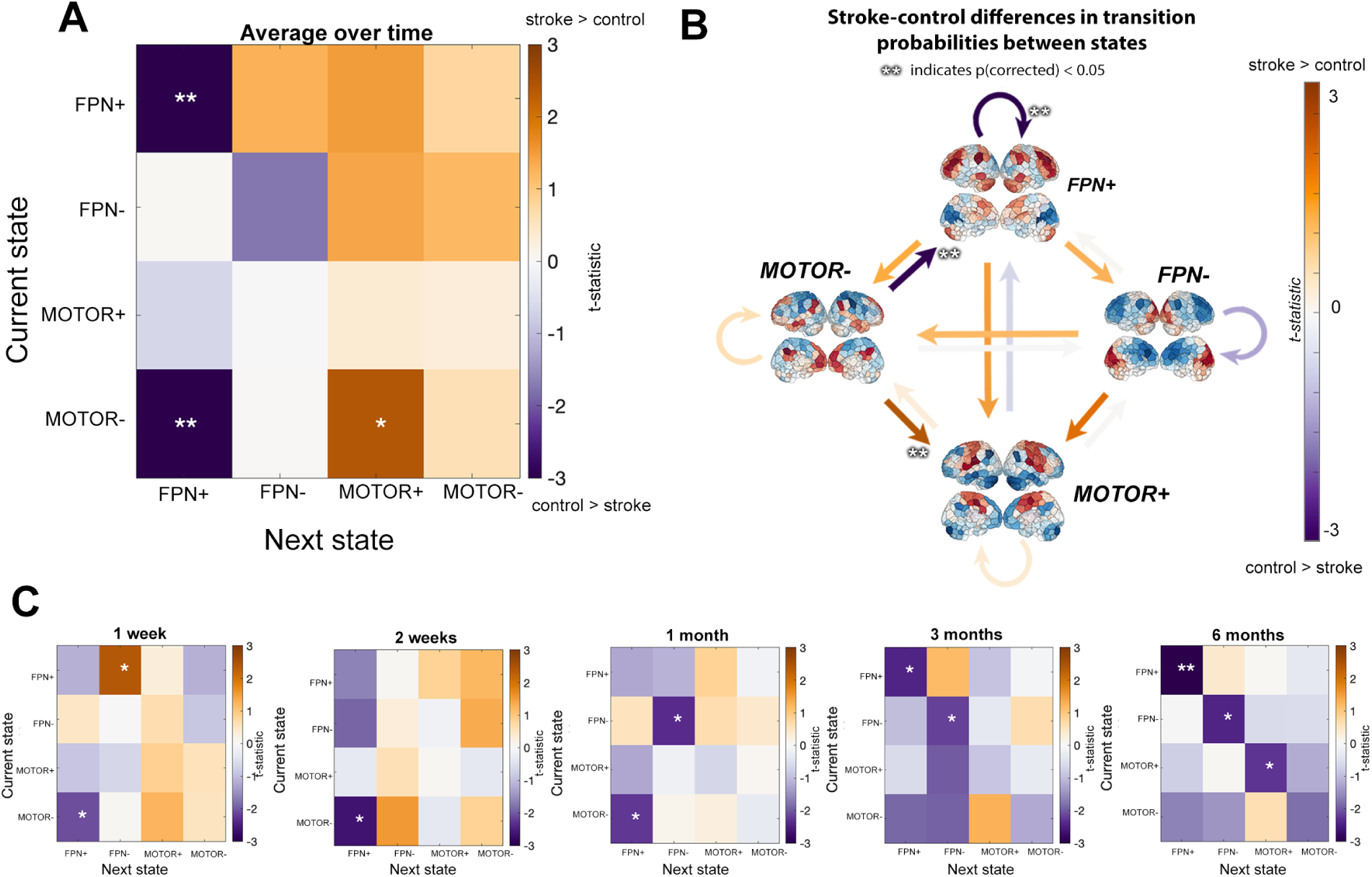
Differences in transition probabilities between stroke and control groups. Transition probabilities include persistence probabilities, i.e. the probability that a state does not transition out of itself. **A**. Differences in average transition and persistence probabilities (over time) between groups. **B**. Same data as panel A, visualized with arrows colored by t-statistic. Double asterisks next to arrows represent a significant group difference that survives multiple comparisons. **C**. Differences in transition probabilities at each follow-up session. T-statistics (stroke-control) are displayed on the color map; double asterisks represent p-values < 0.05 after multiple comparisons correction, single asterisks represent significant uncorrected p-values < 0.05.

### 3.4 Frontoparietal activation relates to motor recovery in individuals with dominant-hand CST damage

At 3 and 6 months post-stroke, we observed a significant association between changes in dwell times of *FPN*^+^ in subjects with dominant-hand CST damage and changes in Fugl-Meyer scores (i.e., the marginal effect 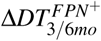), such that increases in (or smaller decreases in) dwell times in these subjects were associated with greater motor improvements (p(FDR) = 0.034 and p(FDR) = 0.034, respectively) (Figure 5B, D). Some individuals with dominant-hand CST damage show decrease in 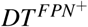 over time, but smaller decreases in 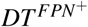 are associated with better recovery. At 3 and 6 months post-stroke, we did not observe a significant association between changes in fractional occupancy of *FPN*^+^ in subjects with dominant-hand CST damage and changes in Fugl-Meyer scores (i.e., the marginal effect 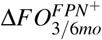) (Figure 5A, C).

**Figure 5:**
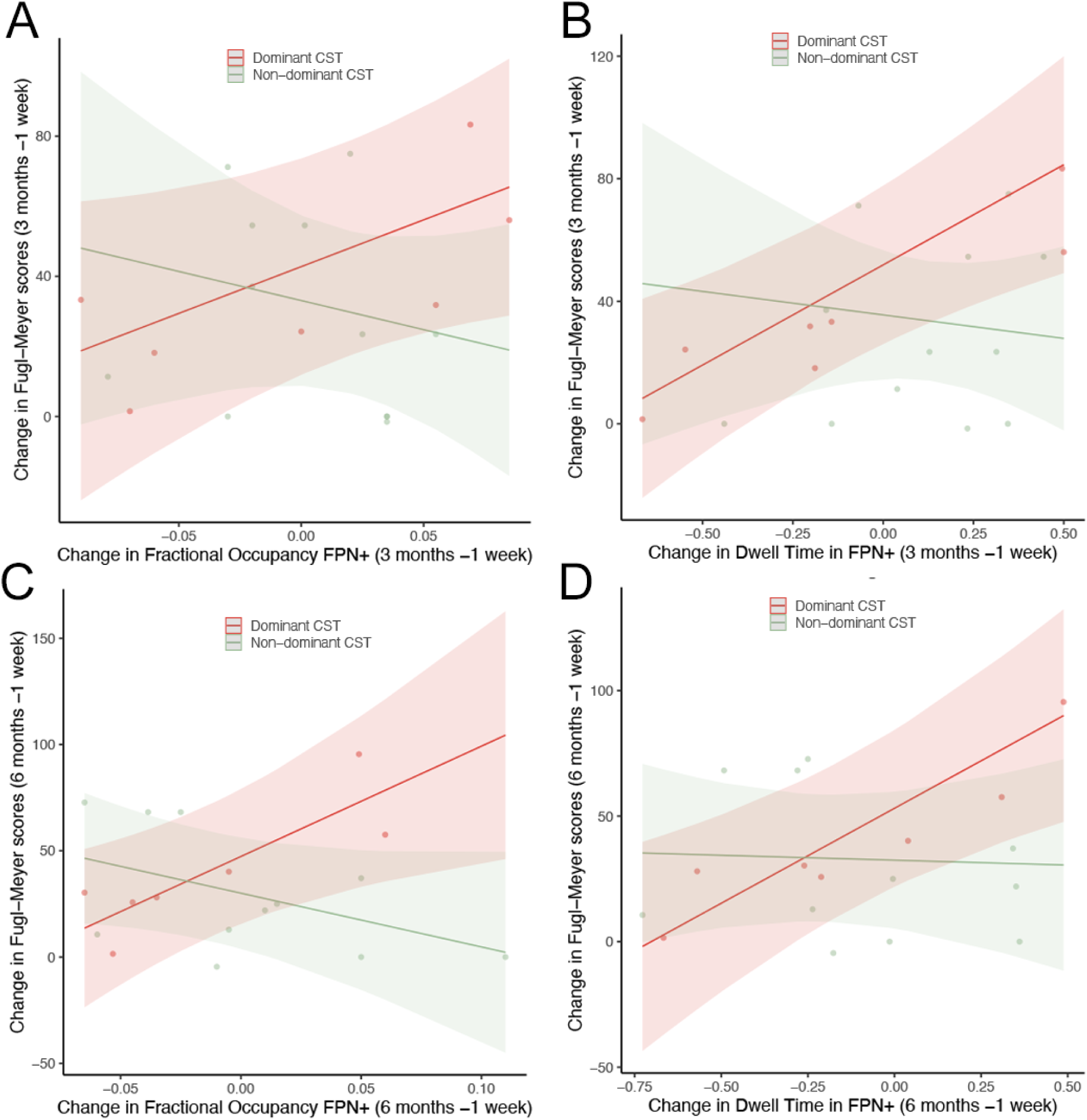
Marginal effects of change in state metrics within the regression model. Marginal effects plots of the change in Fugl-Meyer scores versus the change in fractional occupancy (FO) of *FPN*^+^ from 1 week to 3 months (panel **A**.) and 1 week to 6 months (panel **C**.) post-stroke. Marginal effects plots of the change in Fugl-Meyer scores versus the change in dwell time (DT) of *FPN*^+^ from 1 week to 3 months (panel **B**.) and 1 week to 6 months (panel **D**.) post-stroke.

Finally, we observe that there was a strong and significant correlation between baseline (1 week) FM and 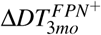 and 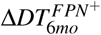 for subjects with dominant-hand CST damage (R = -0.76, p = 0.045, R = -0.83 p = 0.022), see Figure S11A, B. There was not a significant correlation between baseline FM and 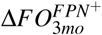 or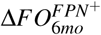 = -0.56, -0.6, p = 0.148, 0.115, respectively). Therefore, we proceeded with creating the null model as described in the Methods section to verify the observed relationships between Δ*FM* and 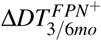 were not a byproduct of the baseline correlation. Indeed, the observed correlation between change in FM and change in DT is always greater than the null distribution of correlation between proportional recovery change in FM and change in DT (p = 0 for 3 and p = 0.01 for 6 months). This gives us confidence that the observed relationships in change in FM and change in DT are not merely due to baseline correlations and proportional recovery.

### 3.5 Fractional occupancy differences are related to functional connectivity differences

Four network pairs had a high magnitude contra-activation (large amplitude activity in opposite directions) in the *FPN*^+^ state: visual I (VIS I) and frontoparietal (FPN), subcortical/cerebellum (SUB) and visual I (VIS I), motor (MOTOR) and subcortical/cerebellum (SUB), and motor (MOTOR) and frontoparietal (FPN). Two pairs of networks had high magnitude co-activation (large amplitude activity in the same direction) in the *FPN*^+^ state: SUB and FPN, and MOTOR and VIS I (Figure 6A). For each individual, we correlated 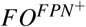 with the FC between each pair co-activated and contra-activated networks, expecting two trends. First, for the highly contra-activated network pairs, we expected a negative correlation between 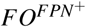 and FC as more time spent in *FPN*^+^ would make the correlation between those networks more negative. Second, for the highly co-activated network pairs, we expected a positive correlation between 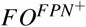 and FC as more time spent in this state would make the correlation between those networks more positive. We did indeed observe the expected relationships for all pairs of networks (Figure 6B). Further, we see that the correlation between 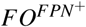 and FC is preserved across and within the groups and is not driven by across-group differences (Simpson’s paradox). Finally, dynamic fluctuations in 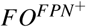 track changes in dynamic FC within frontoparietal areas (Figure 6C, D, E), where the average correlation across subjects between sliding-window FC and 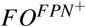 is 0.29 (p = 0, assessed by permutation).

**Figure 6:**
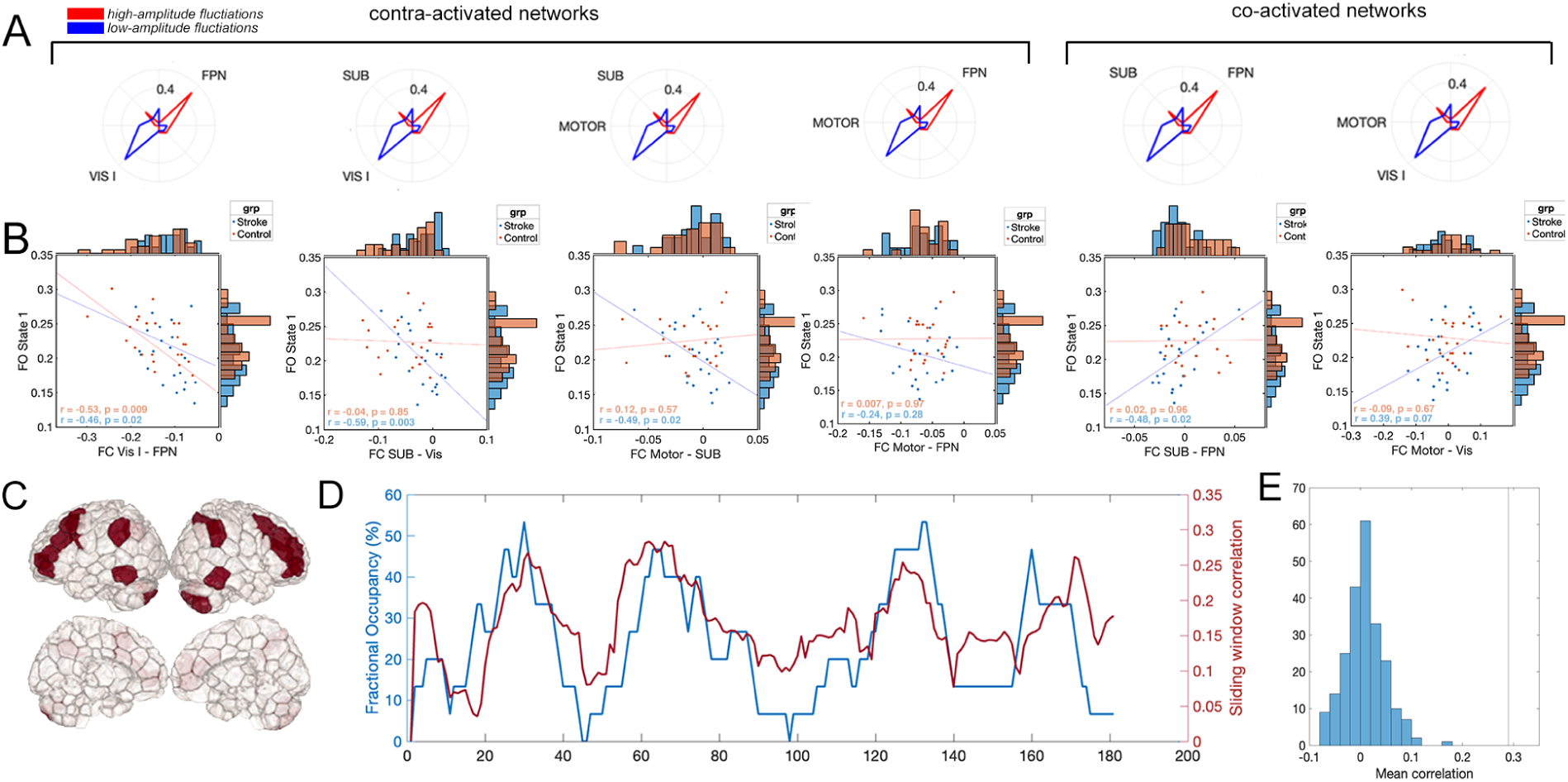
**A**. Pairs of highly activated Yeo networks in *FPN*^+^. **B**. Functional connectivity (FC) between co-activated contra-activated networks in *FPN*^+^ is related to fractional occupancy (FO) of *FPN*^+^ 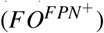 in both stroke and control subjects. Specifically, fewer TRs in a state with strongly contra-activated/co-activated networks results in smaller magnitude FC between those networks. **C**. Areas with high activation in the *FPN*^+^ centroid. **D**. Dynamic, sliding-window fluctuations in 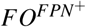 correlate with dynamic FC between regions highly active in the FPN (areas in **C**). **E** Subject-average correlation between dynamic, sliding window 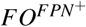 and FC of regions highly active in *FPN*^+^ (black line) and a null distribution of 100 subject-average correlations between dynamic, sliding-window 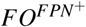 and FC in a set of randomly selected regions (blue histogram).

## 4 Discussion

Large-scale brain activity patterns, or states, can be thought of as group-level temporal building blocks of canonical functional connectivity, reflecting sequences of activity that lie ‘under the hood’ of functional connectivity. Mapping the dynamics of these states offers a fine-grained view into shifts in the temporal sequences of neural activity after stroke that has not yet been explored. In the present study, we provide evidence that spatiotemporal brain dynamics, particularly in a state characterized by high-amplitude frontoparietal activity, are altered after pontine stroke for at least up to 6 months post-infarction. We further demonstrate that increased dwell time in this frontoparietal state is meaningfully related to better improvements in motor function for individuals with dominant-hand CST damage. Finally, we show a direct relationship between time spent in brain states and functional connectivity both in controls as well as individuals with stroke.

### 4.1 Persistent reduced activation of the frontoparietal network in stroke

We observed that stroke subjects spent less time in a brain state characterized by high amplitude frontoparietal network activation, particularly in the sub-acute to chronic stages of stroke (3 and 6 months post-stroke). The frontoparietal network, containing nodes in the frontal cortex like the dorsal premotor cortex and in the parietal cortex like the posterior parietal cortex and intraparietal sulcus, is thought to act as a top-down influence on primary motor networks to control motor output (Marek and Dosenbach, 2018). Communication between nodes in the frontoparietal network is known to be important for computations involved in goal-directed movement, such as motor imagery (Oostra et al., 2016), prospective action judgements (Geers et al., 2021), and generating appropriate hand positions to interact with objects (Borra et al., 2017). Evidence of reduced FC within the frontoparietal network has been observed in subcortical stroke subjects (Wang et al., 2014), as has reduced effective connectivity of the frontoparietal network on the motor network (Inman et al., 2012). Our study extends this current knowledge by demonstrating that the influence of the frontoparietal network in stroke subjects may be weakened, with reduced time spent in states with high amplitude frontoparietal activity.

We did not observe changes in the fractional occupancy, dwell time, or appearance rate of the *MOTOR*^+^ or *MOTOR*^*−*^ states in stroke subjects, as one may anticipate after damage to the corticospinal tract. Instead, we observed changes in the transition probabilities between *MOTOR*^*−*^ and *FPN*^+^, and *MOTOR*^*−*^ and *MOTOR*^+^. These may be more subtle impacts of corticospinal tract disruption that change the sequence of states visited at rest. A lack of differences in the *MOTOR* states could also arise due to the fact that the method cannot detect differences in the magnitude of activations. A critical step in the method involves normalizing each region’s brain activity before clustering. This step would theoretically remove stroke-related differences in the magnitude of activation of motor areas that one may expect after CST damage. As the focus of our paper was to explore stroke-related changes in the temporal patterns of brain activity, we did not explore this possibility, but future work should examine changes in both the magnitude and temporal properties of these states after stroke.

### 4.2 Longer frontoparietal dwell time is associated with better chronic motor recovery in individuals with dominant CST damage

Though reduced frontoparietal network involvement has been observed in stroke subjects relative to controls, as replicated here, there is evidence that within stroke subjects, increased activity of the frontoparietal network is related to better motor recovery (Tscherpel et al., 2020). Recent work has shown that individuals with CST damage (measured using motor-evoked potentials) have greater resting-state functional connectivity in the frontoparietal network compared to individuals without CST damage (Hordacre et al., 2021a). Furthermore, for those subjects with CST damage, better motor recovery was associated with greater resting-state functional connectivity in the frontoparietal network (Hordacre et al., 2021a). We observed that individuals with dominant-hand CST damage had a positive relationship between changes in motor recovery and changes in dwell times in the *FPN*^+^ state from 1 week to 3- and 6-months post-stroke. This result suggests that frontoparietal activation may be an adaptive strategy to support motor recovery which may be more relevant for subjects with dominant hemisphere CST damage.

Increased time spent in the frontoparietal network may be a form of compensation related to structural reserve capacity, a concept believed to reflect increased neural substrate that is neuroprotective against stroke and can modify outcomes (Rosenich et al., 2020). This structural reserve may be particularly well-established to protect against motor impairments in the dominant hemisphere due to the skills and motor programs developed from practice with the dominant hand (Harris and Eng, 2006). Indeed, asymmetries of aspects of frontoparietal tracts have been observed based on handedness, where greater volume in the dominant hemisphere is related to greater voluntary movement planning (Howells et al., 2018), and the residual integrity of frontoparietal fiber tracts is positively associated with motor function in chronic stroke (Schulz et al., 2015). Additionally, descending secondary motor fibers from frontal areas may be able to bypass the CST (Newton et al., 2006) when it is damaged. Thus, it is possible that in subjects with damage to the CST, handedness relative to the lesion side contributes to the reliance on the frontoparietal network in recovery.

### 4.3 Relationship of brain state dynamics and FC

Finally, we showed that across controls and individuals with stroke, FO of the *FPN*^+^ state and FC between networks that dominate that state were related. For example, FC between pairs of networks that were co-activated in the *FPN*^+^ state were positively correlated with FO and FC between pairs of networks that were contra-activated in the *FPN*^+^ state were negatively correlated with FO. Less negative FC between the visual and FPN network was related to less time spent in *FPN*^+^, a state in which the two networks had large magnitude activations in opposite directions. Our findings suggest that time spent in certain brain states may underlie between-group FC differences; a finding which we also replicated with dynamic, sliding window analysis. Knowing how stroke and recovery from stroke is related to time spent in these states may be helpful in designing non-invasive stimulation strategies that are based on direct co-activation/conta-activation of certain regions or networks, not on indirect modulation of FC between pairs of regions/networks.

### 4.4 Clinical relevance

Consideration of state metrics like dwell times, fractional occupancy, appearance rate, and transition probability has the potential to be of great clinical relevance. Stimulation therapies like transcranial magnetic stimulation (TMS) activate or inhibit specific brain areas, often in an attempt to modulate networks whose connectivity has been associated with better recovery (Fisicaro et al., 2019; Grefkes et al., 2010; Fox et al., 2012). These stimulation methods may improve outcomes for those with stroke by permitting the development personalized treatment protocols. Recent TMS modelling work has shown that the effect of regional stimulation on FC depends on the brain state at the time of stimulation (Silvanto and Pascual-Leone, 2008; Edwards et al., 2020). Determining subject-specific metrics of recurring patterns of brain activity may prove to have clinical benefit in the timing and spatial targets of stimulation treatments (Scangos et al., 2021).

Furthermore, the discovery of the effect of the dominant hemisphere in this paper may aid in refining how treatments are individually tailored. In a recent paper, Hordacre et al., 2021b propose a personalized model and suggest targeting alternative brain networks with tDCS. Based on this paper’s findings, we conjecture that this form of stimulation may only be appropriate in those with damage to the dominant hemisphere CST. Additionally, the direction of neural activity changes underlying FC differences across pathological groups compared to control populations has not yet been fully quantified. We showed that analysing FC differences in the context of metrics of brain dynamics provides a more complete picture as to what activation patterns are driving observed differences. If the goal of stimulation therapies is to recapitulate functional network connectivity associated with better outcomes, then understanding how the activations within those networks give rise to connectivity differences may produce more effective targeting strategies.

### 4.5 Limitations

There are several limitations to the study. First, the use of 4 clusters was chosen heuristically, and it is possible that more or fewer brain states exist in the stroke population. Second, k-means-based clustering of the time series does preserve the temporal resolution of brain states (as opposed to static or even dynamic functional connectivity), but with a few caveats. The true activation for a subject at a given time point is most similar to, but not identical to, the discrete cluster centroids described in this paper. There is significant variability in individual activation patterns which may be obscured by concatenating all subjects together and assigning each TR to a single, group-level cluster (Betzel et al., 2021). However, the goal of this study is to derive a meaningful group-level characterization of activity patterns after pontine stroke that may relate to motor recovery. Future work should address extending clustering results to individual subjects in order to support tailored treatments. Additionally, our fMRI sampling rate was 3 seconds which is too slow to capture faster, possibly relevant, brain dynamics (Baker et al., 2014; Kobeleva et al., 2021). Third, the frontoparietal network tends to be lateralized in motor-related activation; however, the clustering approach here uncovered a state with bilateral frontoparietal network activation. Therefore, determining whether contralesional or ipsilesional frontoparietal areas are more involved in recovery in subjects with dominant CST damage was not possible in this analysis. Fourth, the small sample size of subjects with dominant-hand CST damage limits confidence in the findings. Additionally, CST integrity was assessed with the Dice coefficient and a population atlas which may not be accurate for all subject’s neuroanatomy. Finally, most of the individuals with dominant-hand CST damage were right-handed (8/9), and alternative motor functions in the frontal cortex are lateralized to the left premotor cortex (Schluter et al., 2001). Therefore, we cannot fully rule out whether the changes are associated with the left hemisphere or the dominant hemisphere specifically. Similarly, this method is limited in its ability to distinguish the separate contribution of the ipsilateral and contralateral sensorimotor areas, which are known to be differentially activated after stroke (Zemke et al., 2003), as it seeks to determine recurring states present in healthy controls and stroke subjects. As a result, we expect that our results reflect adaptations/disruptions to “canonical” brain activation states present across stroke and control populations (and may change after stroke in the proportion or frequency with which they are expressed). Further work that investigates more states, possibly states specific to the post-CST stroke brain, may elucidate the role of contralesional hemisphere activation, but that is beyond the scope of this study.

### 4.6 Conclusions

This work suggests that when the dominant-hand CST is compromised in stroke, resting-state configurations may shift to include increased activation of alternative network configurations, like frontoparietal regions, to promote compensatory neural pathways that support recovery of motor function when traditional motor circuits of the dominant-hemisphere CST are compromised. Determining subject- and stroke lesion-specific patterns of brain activity highly associated with recovery may help improve the design of clinical rehabilitation strategies, including non-invasive brain stimulation.

---

**Figure S1:**
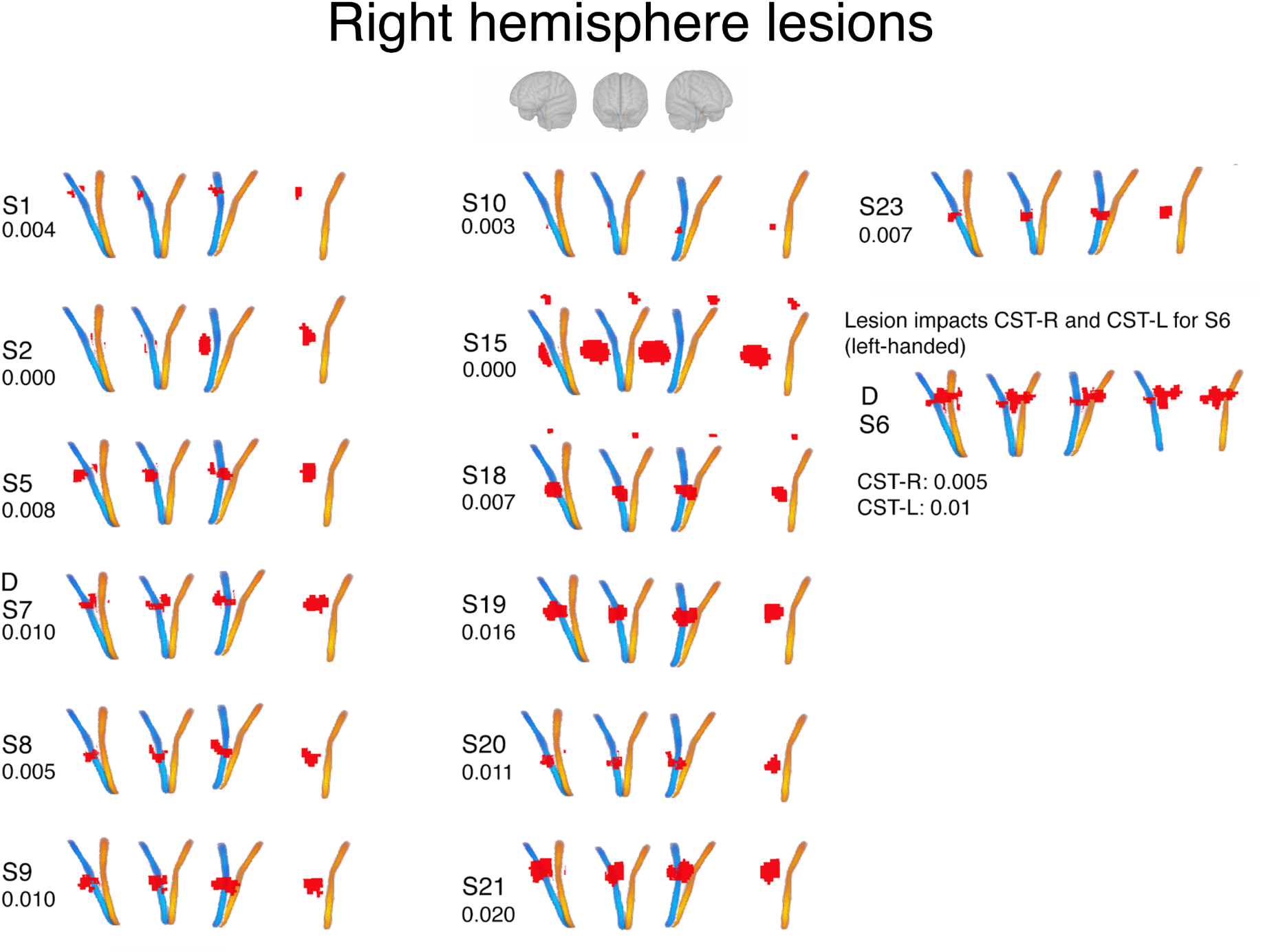
Right hemisphere lesion location relative to brainstem corticospinal tracts. Red = lesion. Blue = right CST, yellow = left CST. Four views of the lesions/CSTs are displayed, from left to right: left lateral view, anterior view, right lateral view. A “D” above the subject identifiers (“SX”) indicates that the lesion is in that subject’s dominant hemisphere. Numbers below subject identifiers indicate Dice overlap between lesion and ipsilesional CST.

## Supplementary Figures

**Figure S2:**
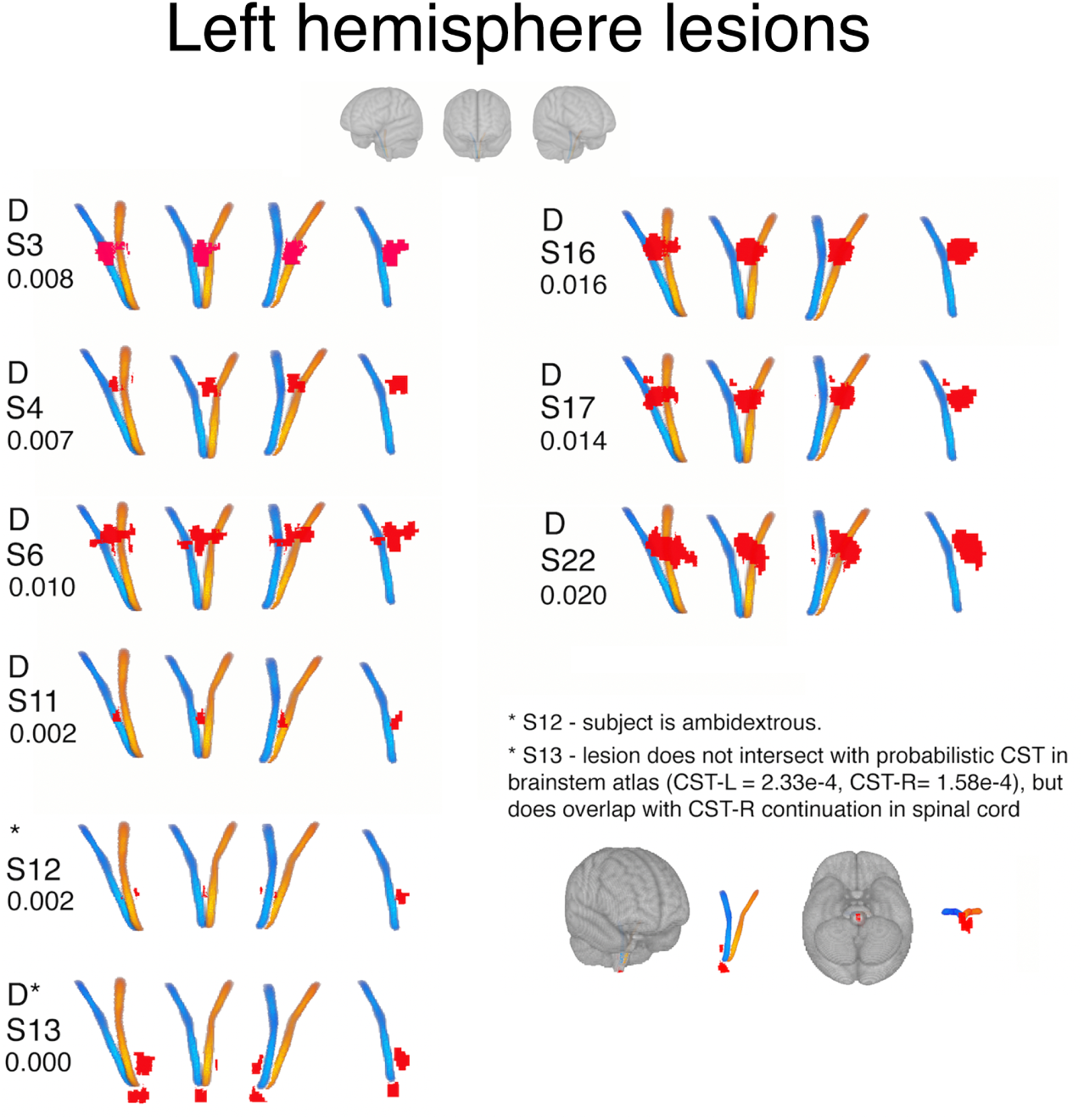
Left hemisphere lesion location relative to brainstem corticospinal tracts. Red = lesion. Blue = right CST, yellow = left CST. Four views of the lesions/CSTs are displayed, from left to right: left lateral view, anterior view, right lateral view. A “D” above the subject identifiers (“SX”) indicates that the lesion is in that subject’s dominant hemisphere. Numbers below subject identifiers indicate Dice overlap between lesion and ipsilesional CST.

**Figure S3:**
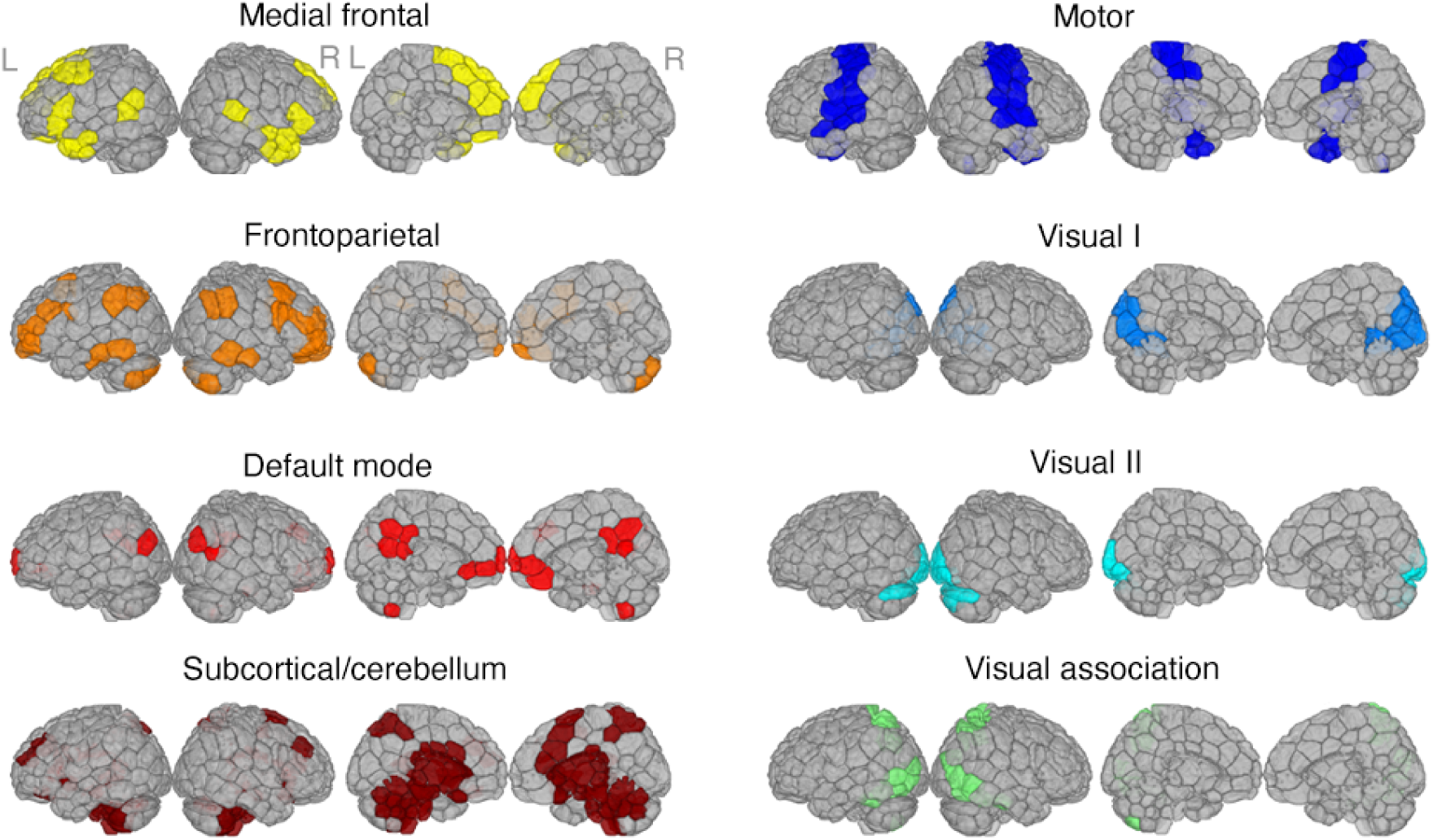
Functional networks used in analysis (based on Shen et al., 2013) that represent communities of brain regions with similar activity over time.

**Figure S4:**
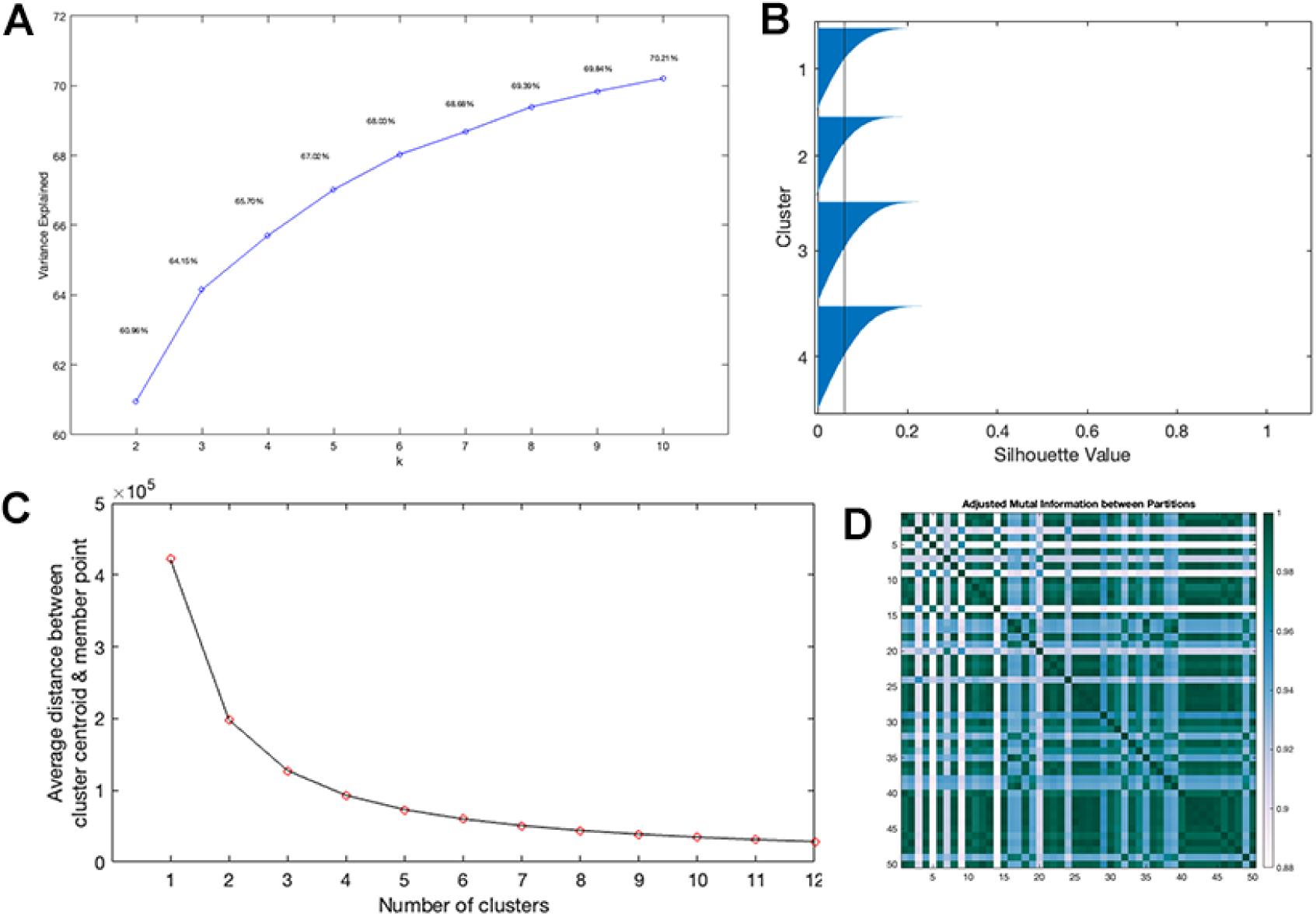
Clustering criteria used to determine the optimal cluster number. **A**. Elbow plot: the total variance explained by increasing k from from k=3 to k = 4 is 1.4%; from k = 4 to k = 5 is 1.2%. **B**. Silhouette values for k = 4 where negative values indicate that a data point (TR) may have been assigned to the wrong cluster. **C**. Average distance between cluster centroids and each member point (distortion criteria). **D**. Adjusted mutual information between 50 final clusters with varied initialization of k-means when k is set to 4 (minimum value observed = 0.88).

**Figure S5:**
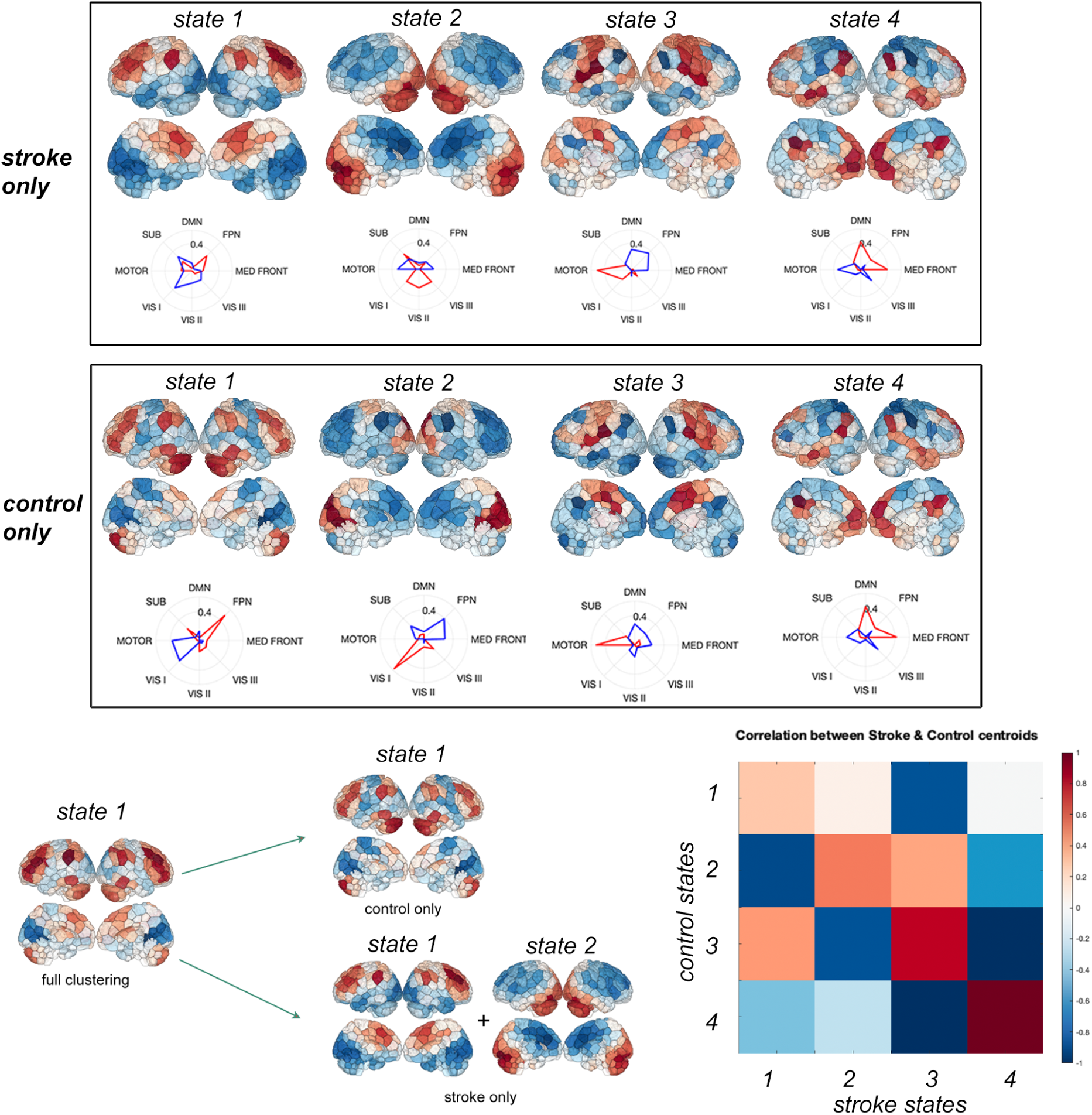
Centroids derived from clustering stroke and control subjects separately. Bottom left: Hypothesized mapping from the centroids derived from clustering both groups together to the individually clustered centroids. Bottom right: Correlation of the centroids derived from separately clustering control and stroke groups.

**Figure S6:**
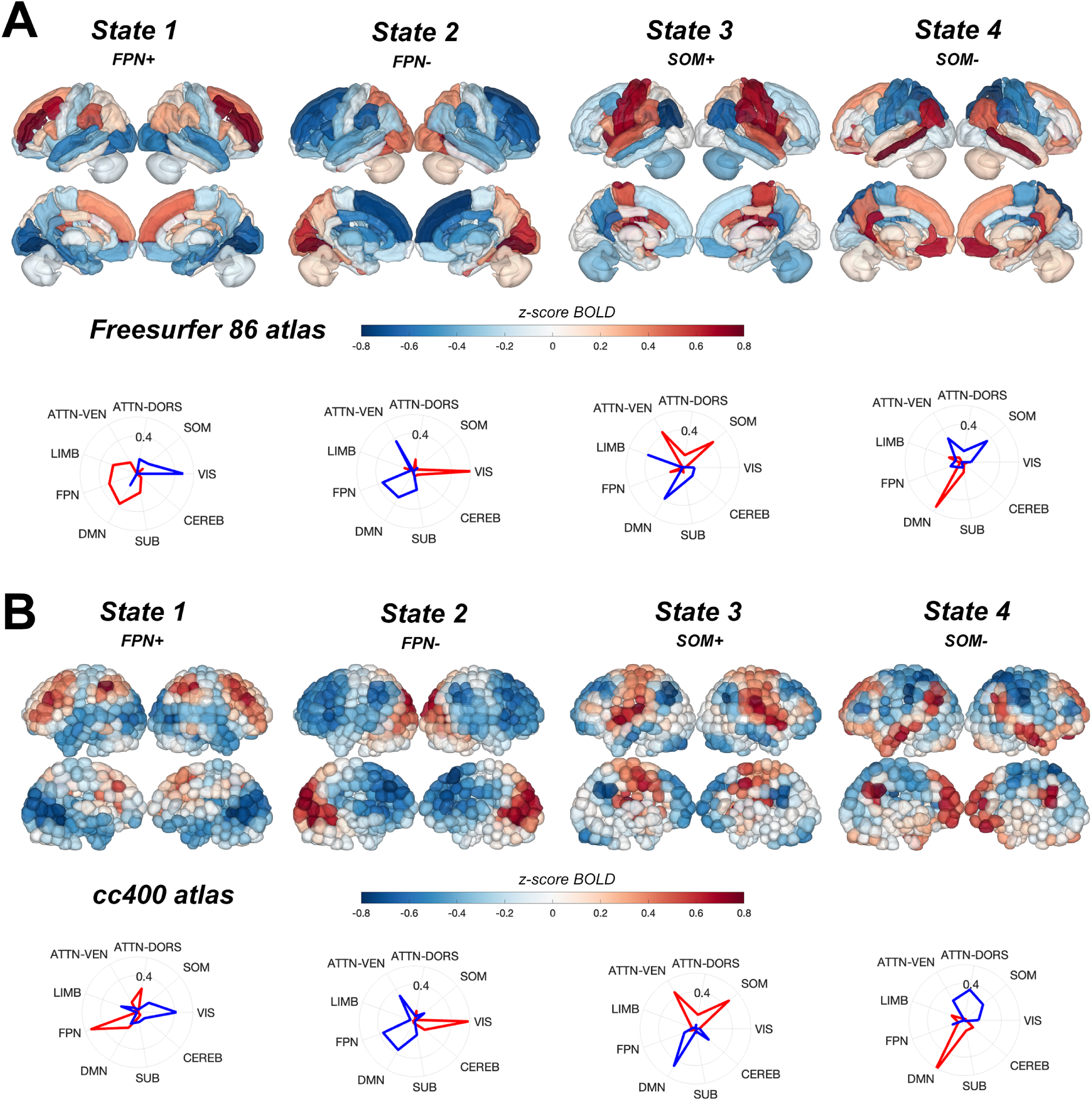
Centroids derived from clustering fMRI data parcellated using different atlases: **A** FreeSurfer-86 region (group-average; not individual anatomical parcellations) and **B** CC400. Note that the regions are parcellated into 9 networks here, instead of 8, and there are missing (VIS II, VIS III, MED FRONT) and additional (ATTN-VEN, ATTN-DORS, LIMB) networks that are not in the shen268 network parcellation. SUB + CEREB are also combined in the shen268 parcellation, and the label SOM is used here instead of MOTOR in the main results.

**Figure S7:**
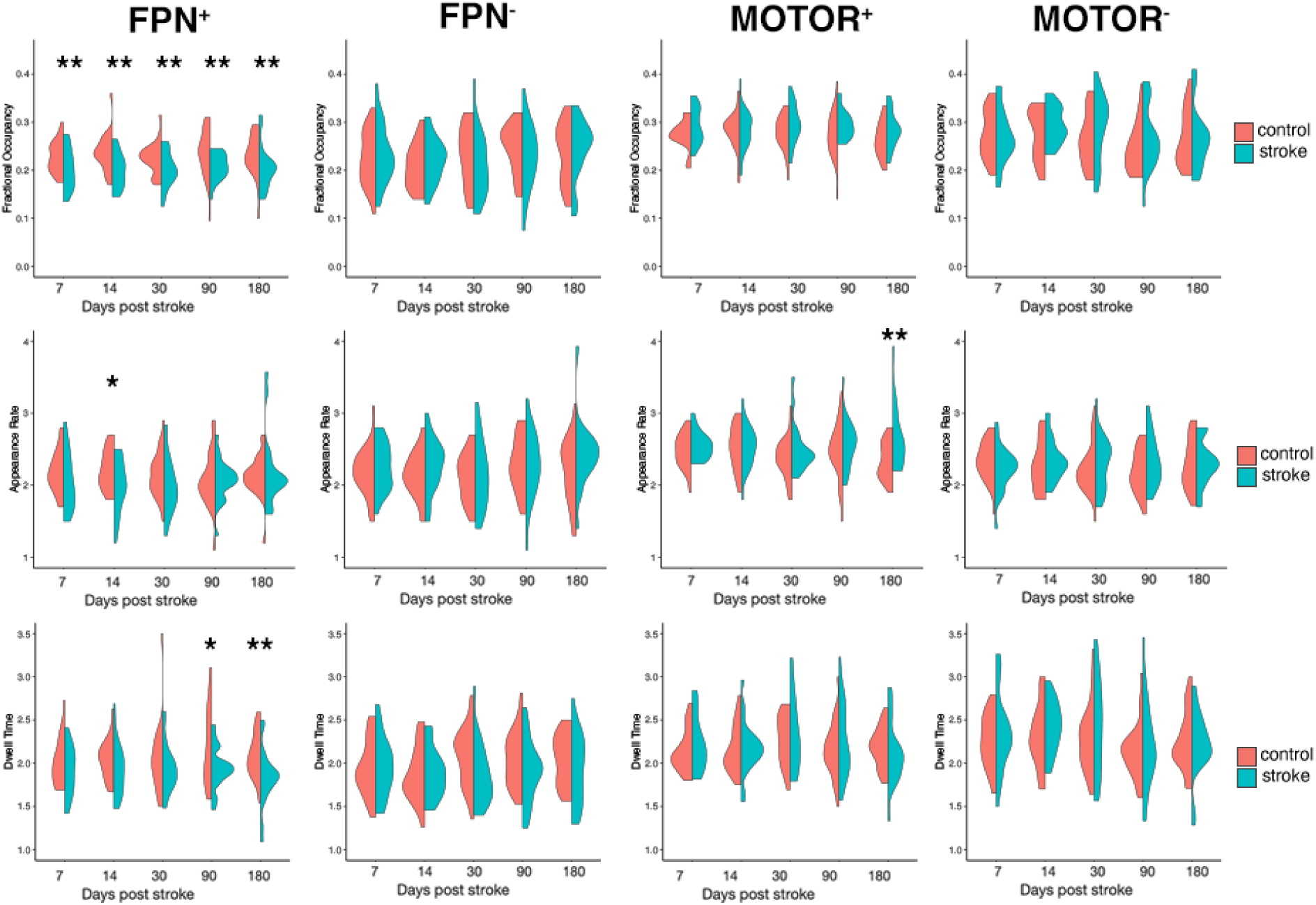
Full session-specific group differences in fractional occupancy (top row), dwell time (middle row), and appearance rate (bottom romw) with k = 4. Aside from the group differences reported in the main paper in *FPN*^+^, there are group differences in 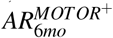.

**Figure S8:**
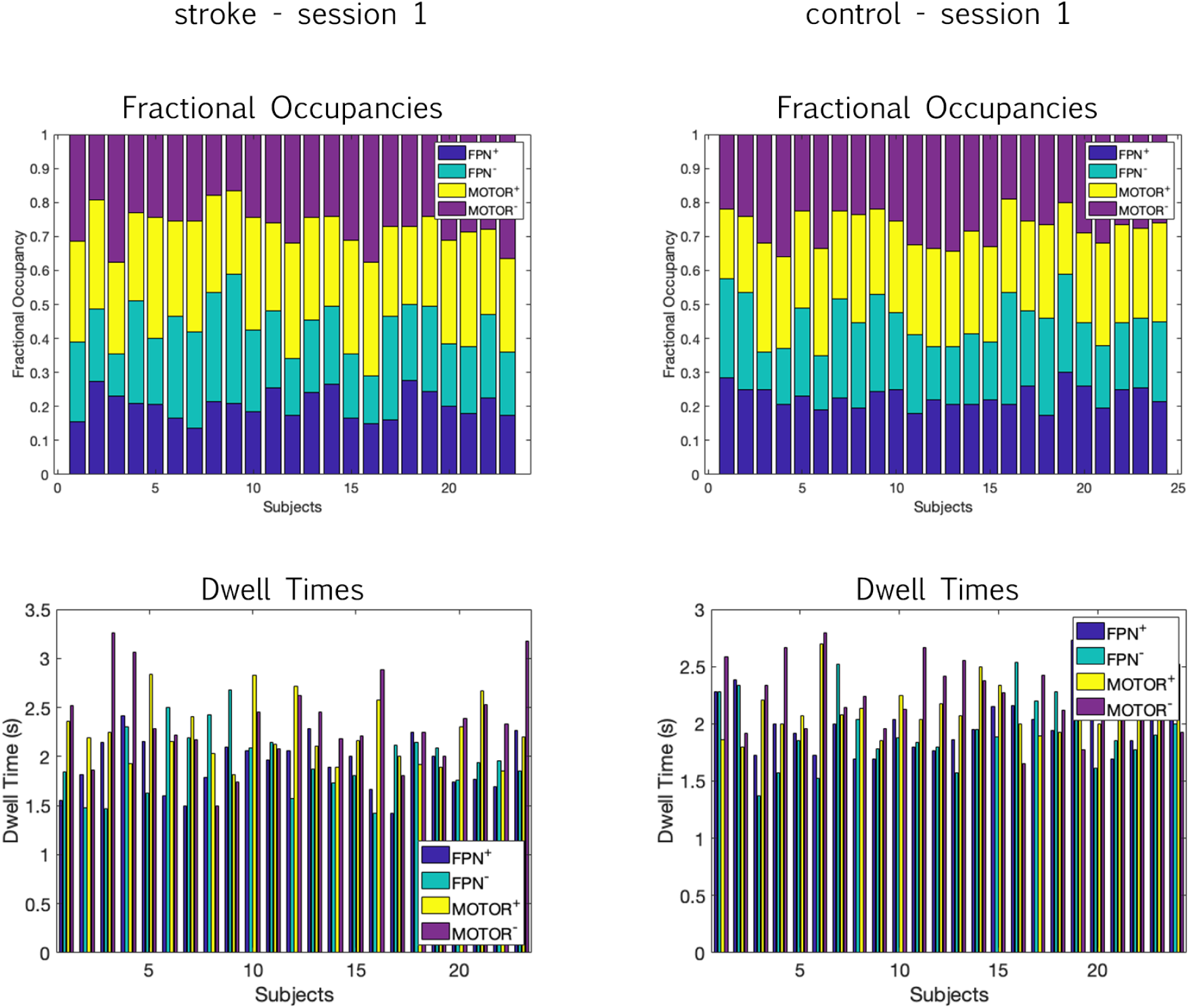
Dwell times and fractional occupancies of stroke subjects (left) and control subjects (right) at session 1.

**Figure S9:**
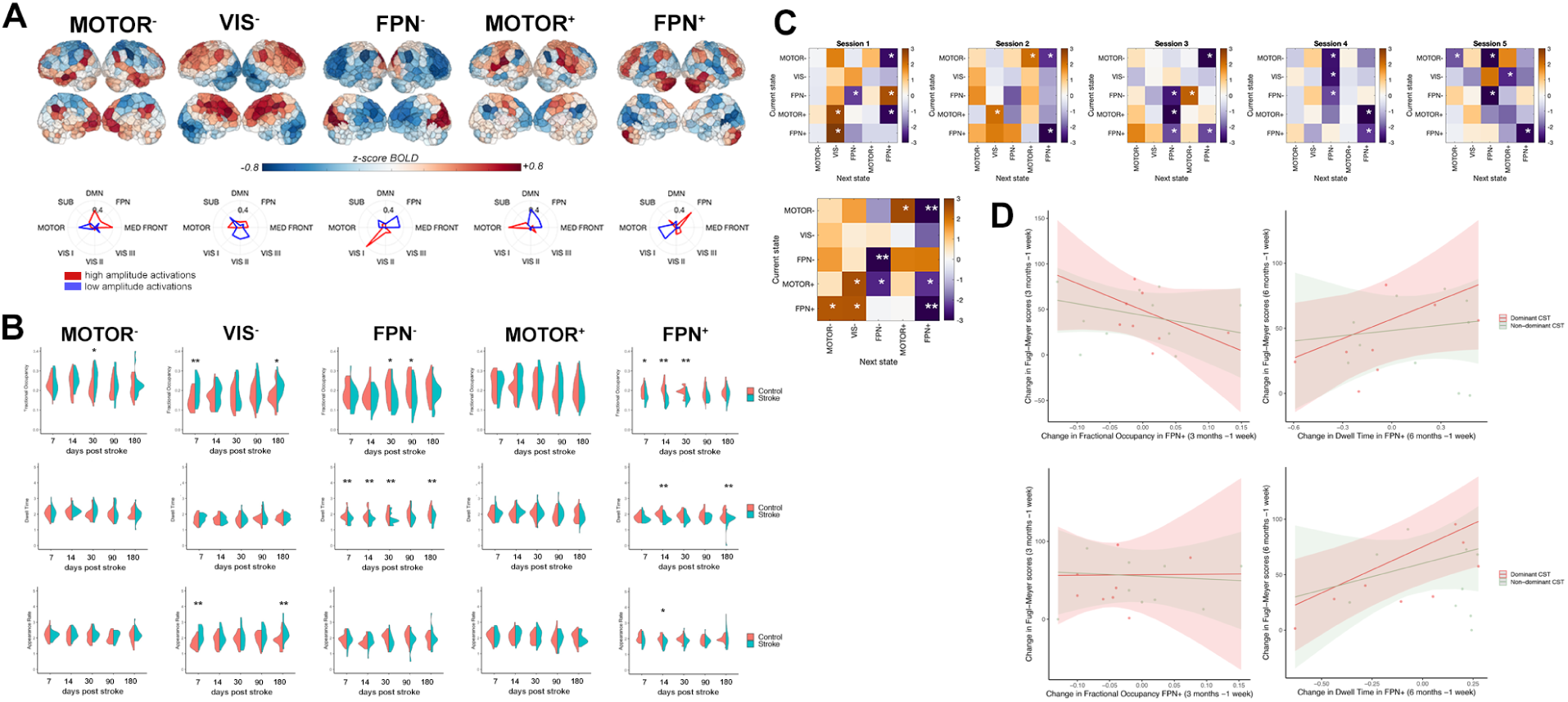
Main results from the paper replicated with k = 5 clusters. **A**. Cluster centroids from k = 5 show that largely the same states appear (*FPN*^+^,*FPN*^*−*^, *MOTOR*^+^, *MOTOR*^*−*^) alongside a new state (*V IS*^*−*^) characterized by low amplitude activations of the visual network. **B**. Stroke-control differences in fractional occupancy (top row), dwell time (middle row), and appearance rate (bottom row) mirror differences observed with k = 4, particularly group differences in FO and DT in *FPN*^+^. Notably, *FPN*^*−*^ displays significant group differences in DT using k = 5 but not k = 4. **C**. Stroke-control differences in transition probabilities between states mirrors results with k = 4, particularly reduced transition probability from *MOTOR*^*−*^ into *FPN*^+^ and reduced persistence probability of *FPN*^+^. **D**. Linear model results examining the relationship between longitudinal changes in 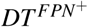 and 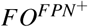 and motor recovery in subjects with dominant hemisphere CST damage. Trend-level effects are replicated with k = 5 but do not reach statistical significance (p-value of marginal effect of 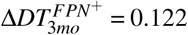 (uncorrected), 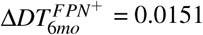 (uncorrected).

**Figure S10:**
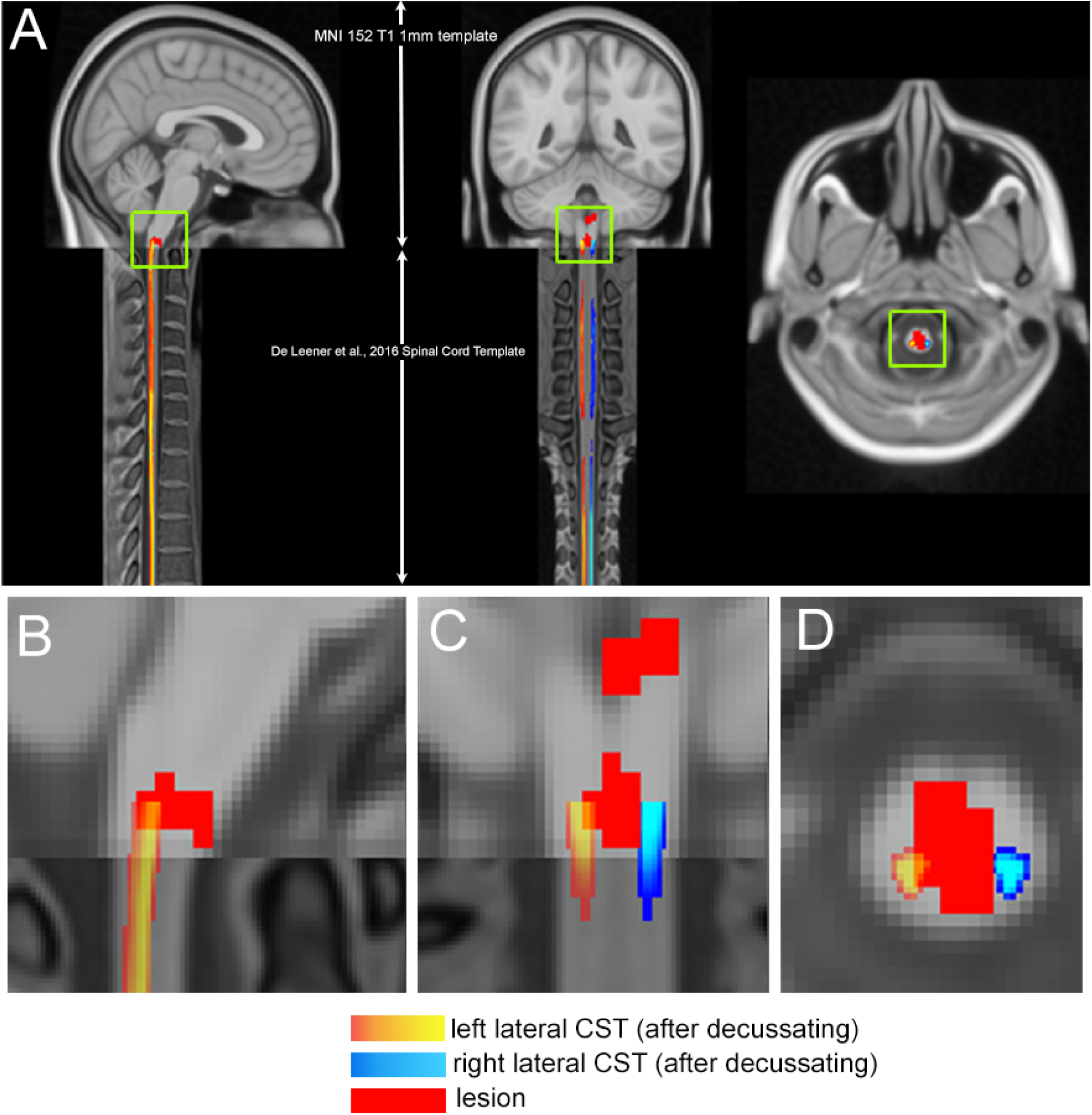
A.Sagittal, coronal, and horizontal views of the lateral corticospinal tract atlases and lesion for subject S13 (who is right-handed) on top of the MNI 152 brain template (top) and De Leneer et al. spinal cord template. Close-ups of B., sagittal slice, C., coronal slice, and D., horizontal slice with reduced opacity of the left lateral CST, which is the dominant CST as the subject is right-handed and the lesion is in the right spinal cord after decussating in the medulla.

**Figure S11:**
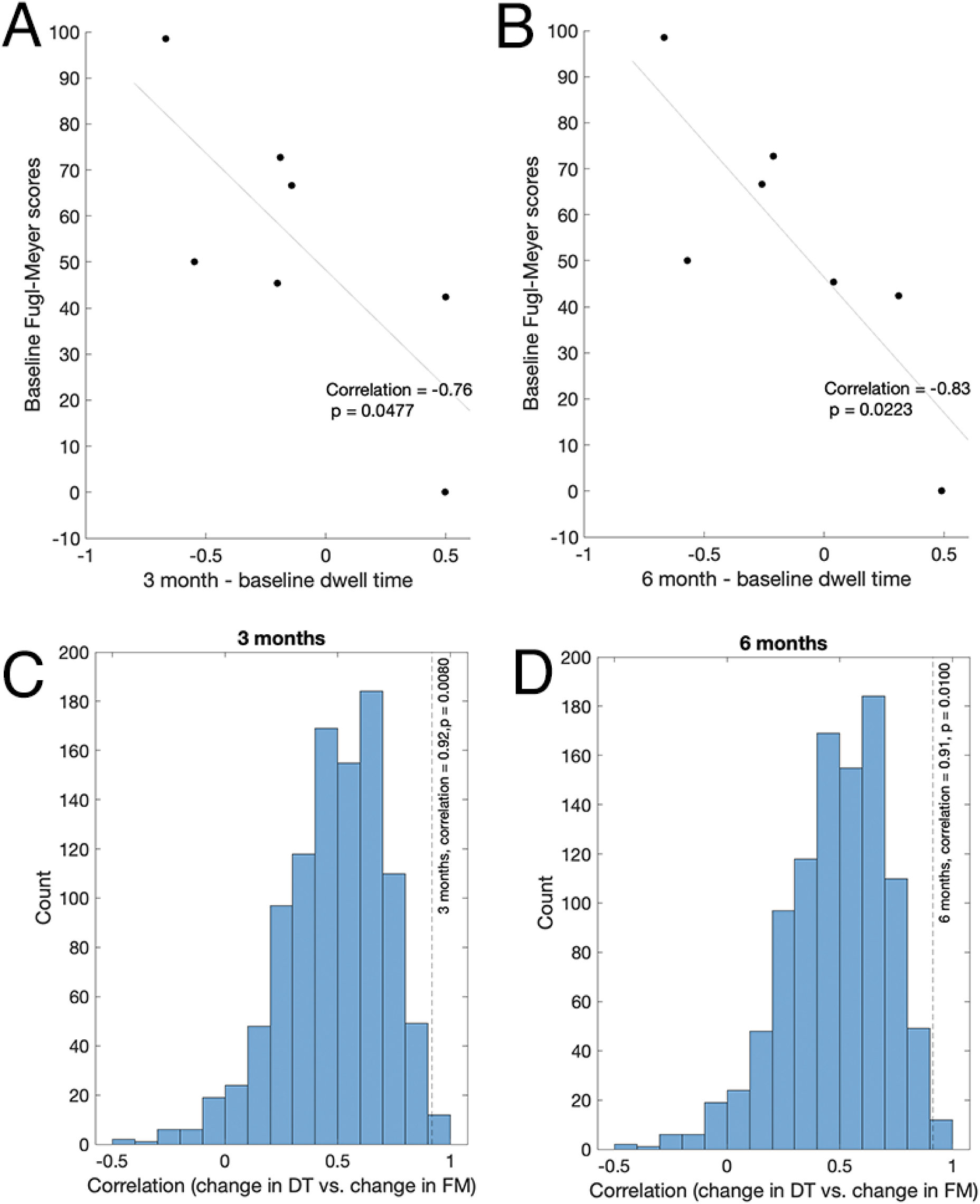
**A**. Correlation between 1-week (baseline) Fugl-Meyer scores, and the change in dwell time in *FPN*^+^ between 1 week and 3 months in subjects with dominant hemisphere CST damage. **B**. Correlation between 1-week (baseline) Fugl-Meyer scores, and the change in dwell time in *FPN*^+^ between 1 week and 6 months in subjects with dominant hemisphere CST damage. **C**. Distribution of correlations between change in dwell time and change in Fugl-Meyer using observed 1-week and 3-month dwell times, observed 1 week Fugl-Meyer scores, and 3-month Fugl-Meyer scores created according to the proportional recovery plus random noise (100 sets of 3-month Fugl-Meyer scores were generated). Observed correlation using observed 3 month Fugl-Meyer scores is shown as the grey dotted line (R = 0.92). **D**. Distribution of correlations between change in dwell time and change in Fugl-Meyer using observed 1-week and 6-month dwell times, observed 1 week Fugl-Meyer scores, and 6-month Fugl-Meyer scores created according to the proportional recovery plus random noise (100 sets of 6-month Fugl-Meyer scores were generated). Observed correlation using observed 6 month Fugl-Meyer scores is shown as the grey dotted line (R = 0.91).

**Figure S12:**
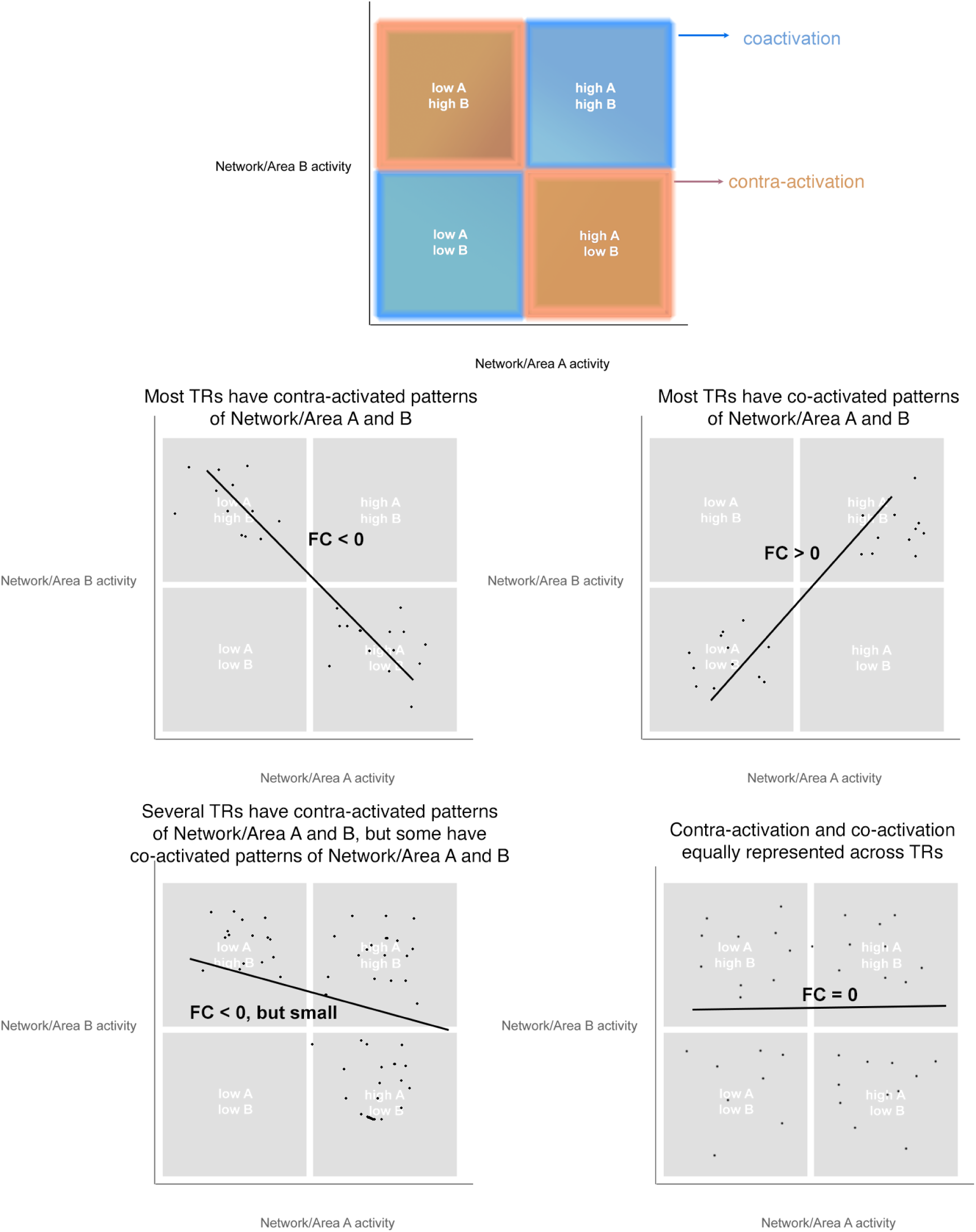
Theoretical framework for linking fractional occupancy of brain states to functional connectivity between brain regions. Each TR can be plotted into one of four quadrants representing the joint activity patterns of a given pair of networks: high activity of networks/regions A and B (coactivation); low activity of A and B (coactivation); high A, low B (contra-activation); and low A, high B (contra-activation). When most TRs are in a contra-activation configuration of 2 areas, then the functional connectivity between those areas will be negative; when most TRs are in a co-activation configuration, the FC between those two areas will be positive.

## Notes

### Competing Interest Statement

The authors have declared no competing interest.

