## Supplementary Figures for "Increased prevalence of a frontoparietal brain state is associated with better motor recovery after stroke affecting dominant-hand corticospinal tract"

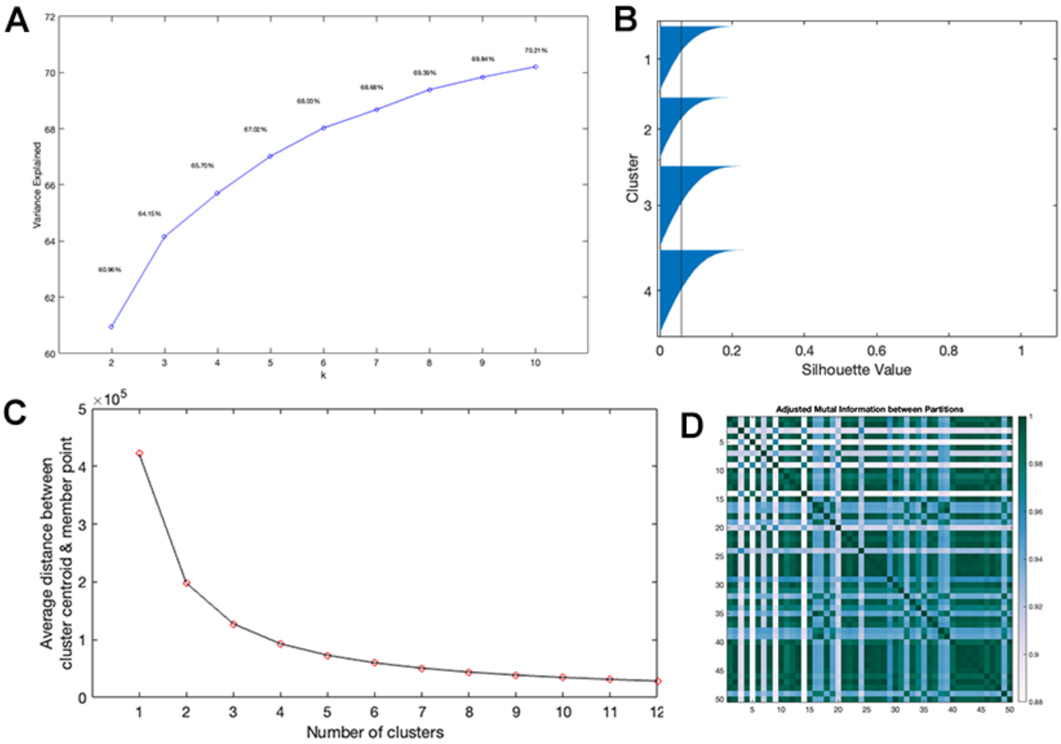


Figure S1: Clustering criteria used to determine the optimal cluster number. **A.** Elbow plot: the total variance explained by increasing k from from k=3 to k = 4 is 1.4%; from k = 4 to k = 5 is 1.2%. **B.** Silhouette values for k = 4 where negative values indicate that a data point (TR) may have been assigned to the wrong cluster. **C.** Average distance between cluster centroids and each member point (distortion criteria). **D.** Adjusted mutual information between 50 final clusters with varied initialization of k-means when k is set to 4 (minimum value observed = 0.88).

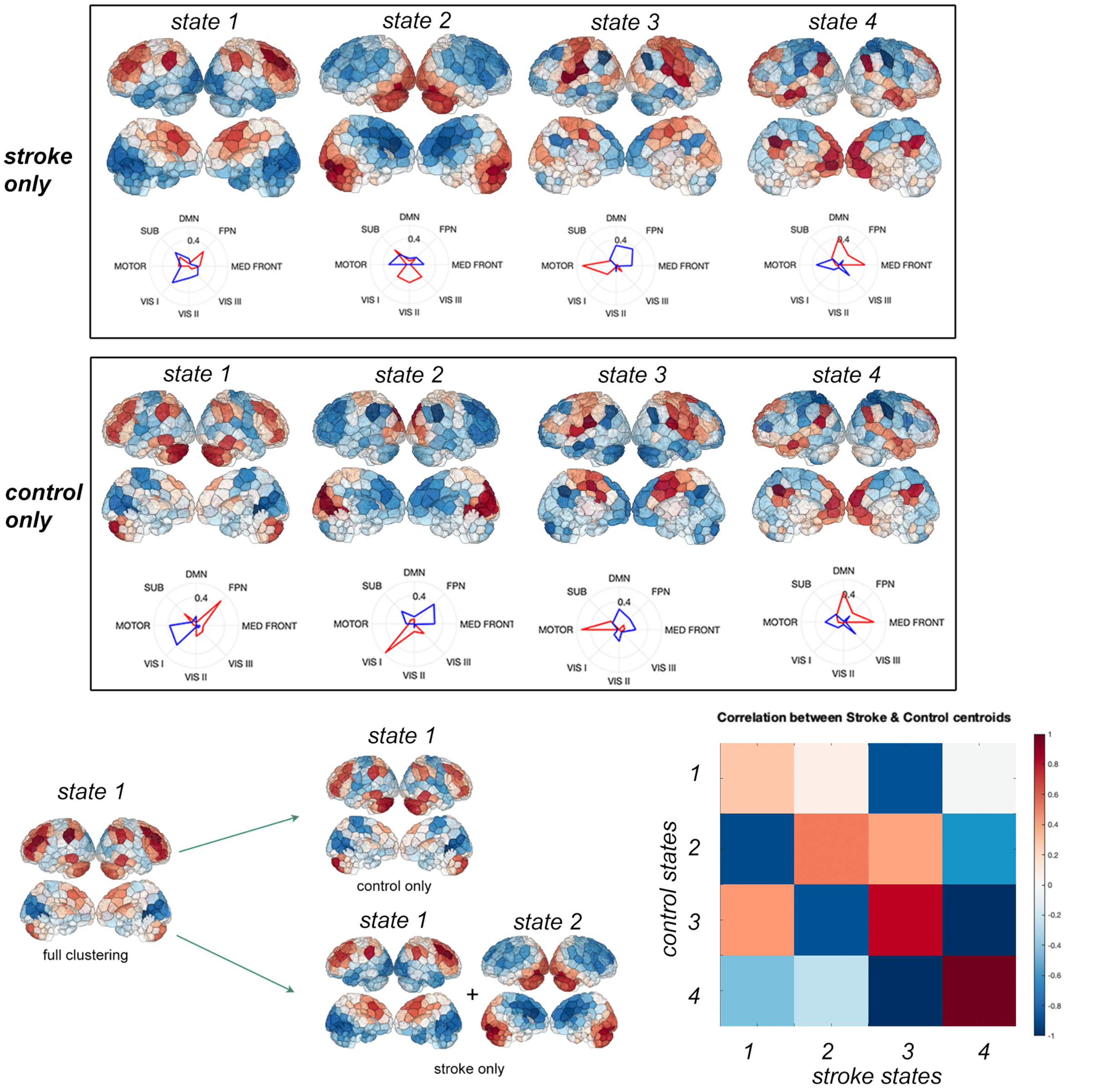


Figure S2: Centroids derived from clustering stroke and control subjects separately. Bottom left: Hypothesized mapping from the centroids derived from clustering both groups together to the individually clustered centroids. Bottom right: Correlation of the centroids derived from separately clustering control and stroke groups.

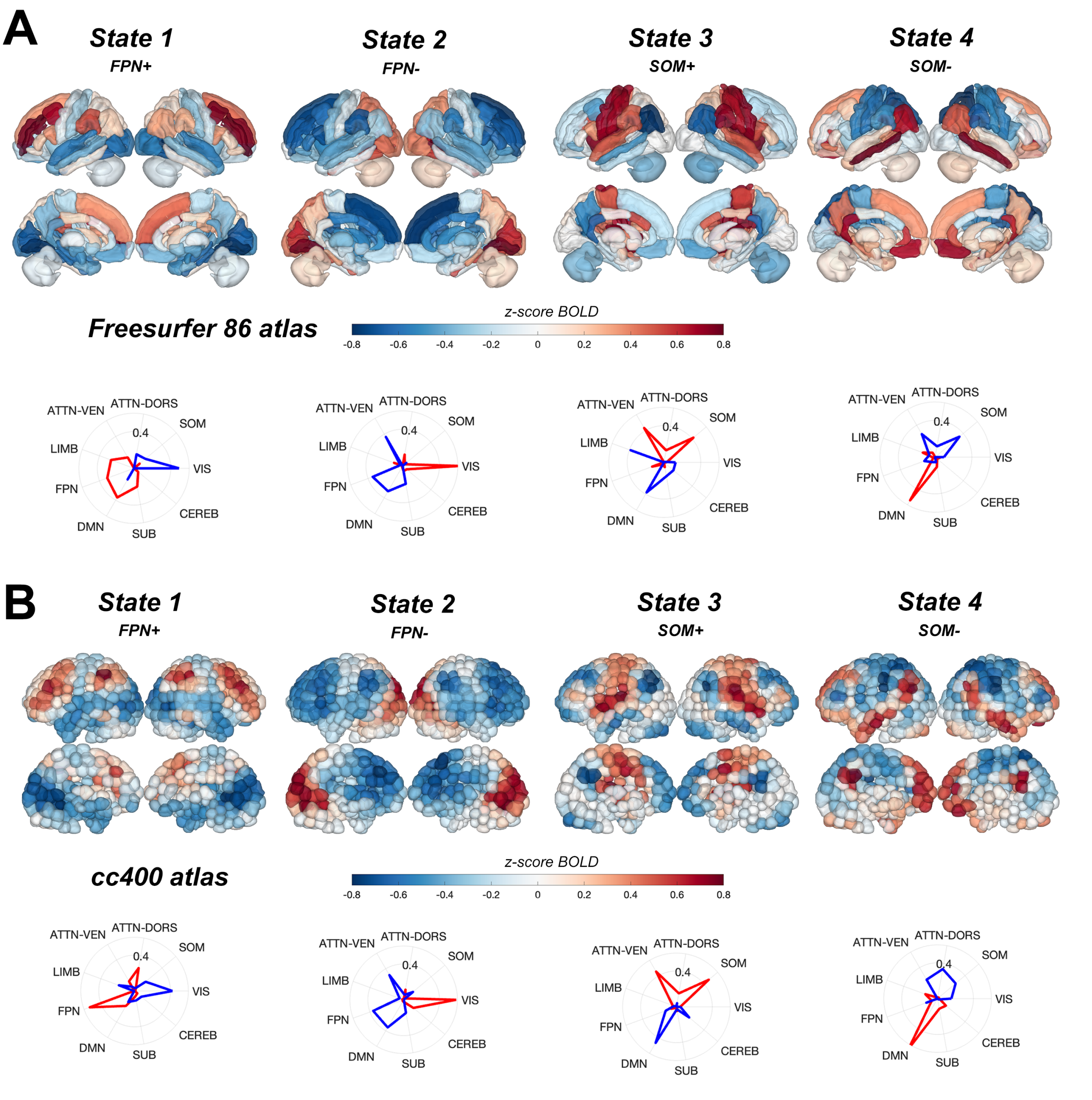


Figure S3: Centroids derived from clustering fMRI data parcellated using different atlases: **A** FreeSurfer-86 region (group-average; not individual anatomical parcellations) and **B** CC400 . Note that the regions are parcellated into 9 Yeo networks here, instead of 8, and there are missing (VIS II, VIS III, MED FRONT) and additional (ATTN-VEN, ATTN-DORS, LIMB) networks that are not in the shen268 Yeo parcellation. SUB + CEREB are also combined in the shen268 parcellation, and the label SOM is used here instead of MOTOR in the main results.

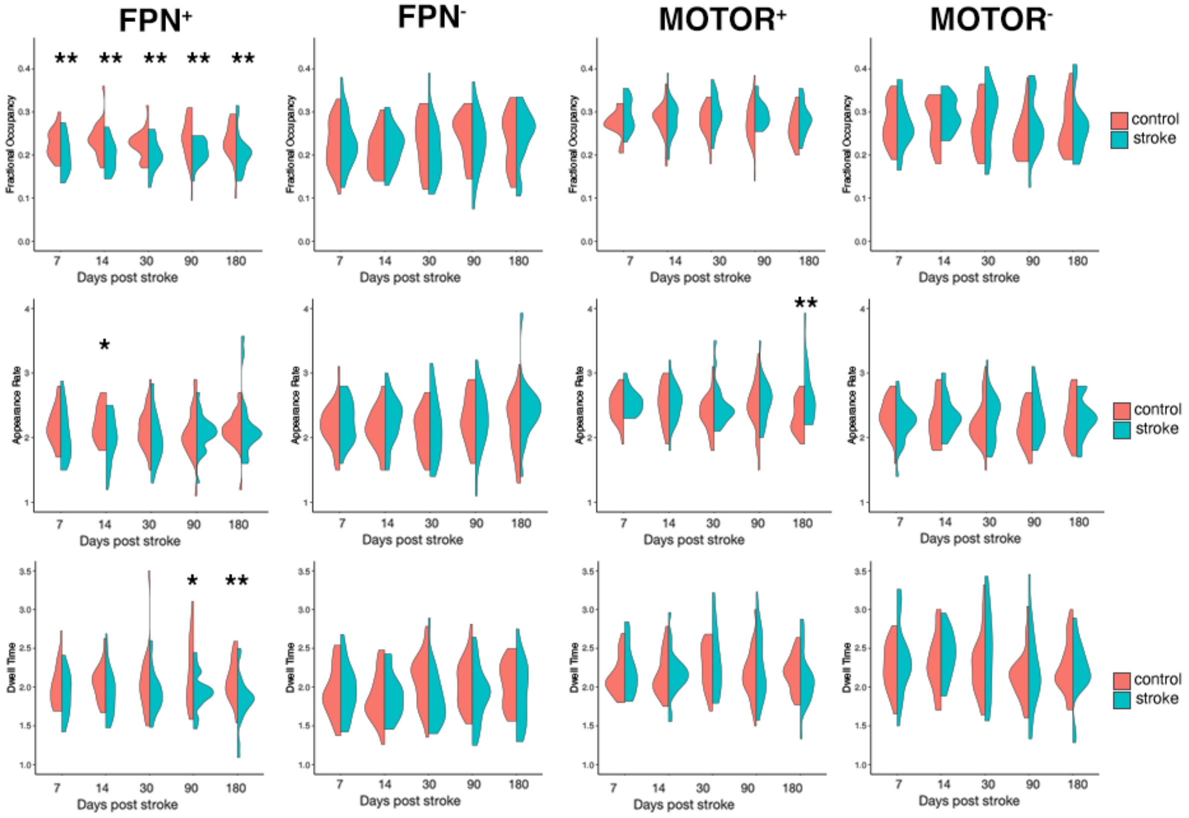


Figure S4: Full session-specific group differences in fractional occupancy (top row), dwell time (middle row), and appearance rate (bottom romw) with k = 4. Aside from the group differences reported in the main paper in $FPN^{+}$, there are group differences in $AR_{6mo}^{MOTOR^{+}}$.

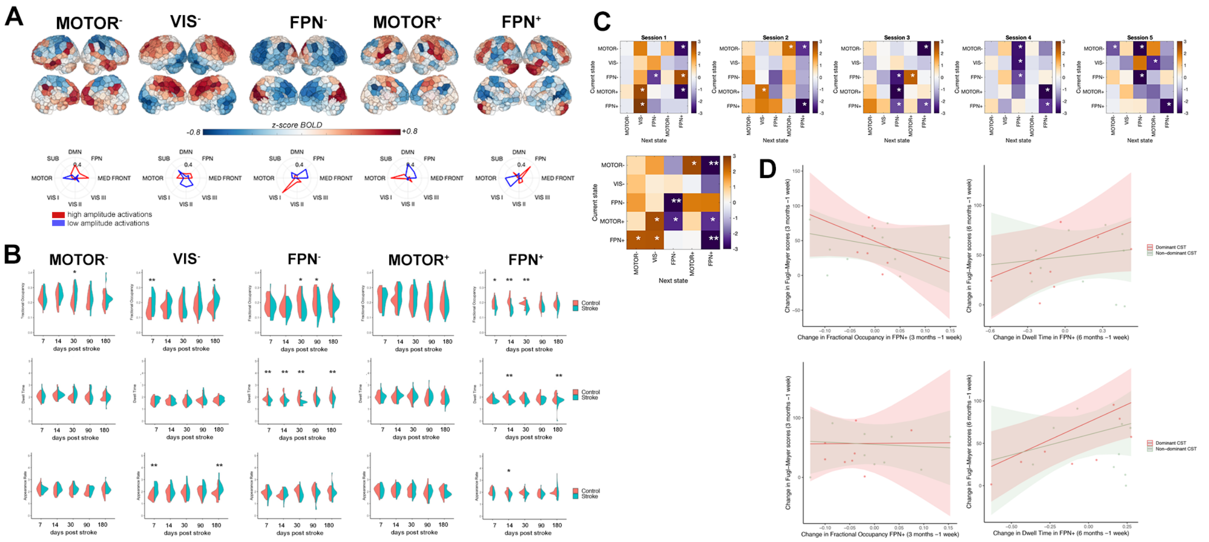


Figure S5: Main results from the paper replicated with k = 5 clusters. **A**. Cluster centroids from k = 5 show that largely the same states appear ($FPN^{+}$,$FPN^{-}$, $MOTOR^{+}$, $MOTOR^{-}$) alongside a new state ($VIS^{-}$) characterized by low amplitude activations of the visual network. **B**. Stroke-control differences in fractional occupancy (top row), dwell time (middle row), and appearance rate (bottom row) mirror differences observed with k = 4, particularly group differences in FO and DT in $FPN^{+}$. Notably, $FPN^{-}$ displays significant group differences in DT using k = 5 but not k = 4. **C.** Stroke-control differences in transition probabilities between states mirrors results with k = 4, particularly reduced transition probability from $MOTOR^{-}$ into $FPN^{+}$ and reduced persistence probability of $FPN^{+}$. **D**. Linear model results examining the relationship between longitudinal changes in $DT^{FPN^{+}}$ and $FO^{FPN^{+}}$ and motor recovery in subjects with dominant hemisphere CST damage. Trend-level effects are replicated with k = 5 but do not reach statistical significance (p-value of marginal effect of $\Delta DT_{3mo}^{FPN^{+}}$ = 0.122 (uncorrected), $\Delta DT_{6mo}^{FPN^{+}}$ = 0.0151 (uncorrected).

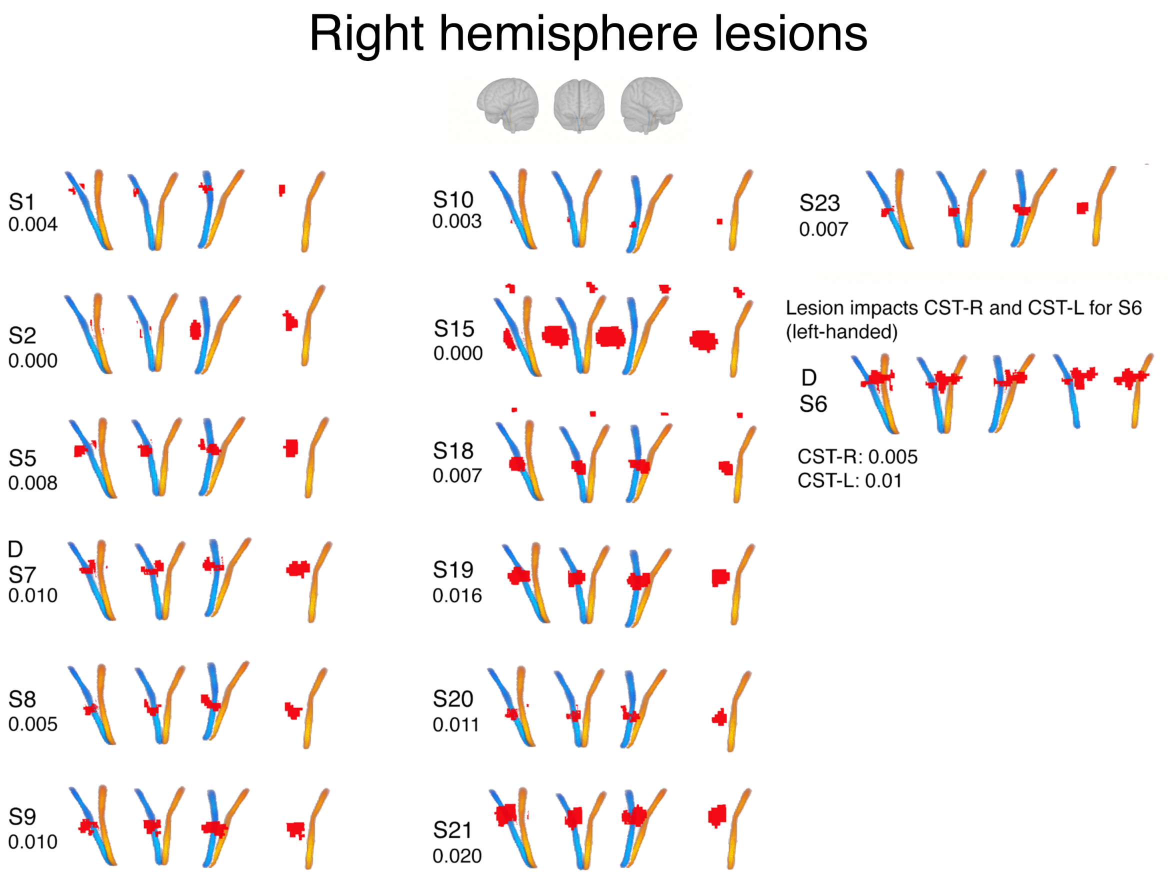


Figure S6: Right hemisphere lesion location relative to brainstem corticospinal tracts. Red = lesion. Blue = right CST, yellow = left CST. Four views of the lesions/CSTs are displayed, from left to right: left lateral view, anterior view, right lateral view. A "D" above the subject identifiers ("SX") indicates that the lesion is in that subject's dominant hemisphere. Numbers below subject identifiers indicate Dice overlap between lesion and ipsilesional CST.

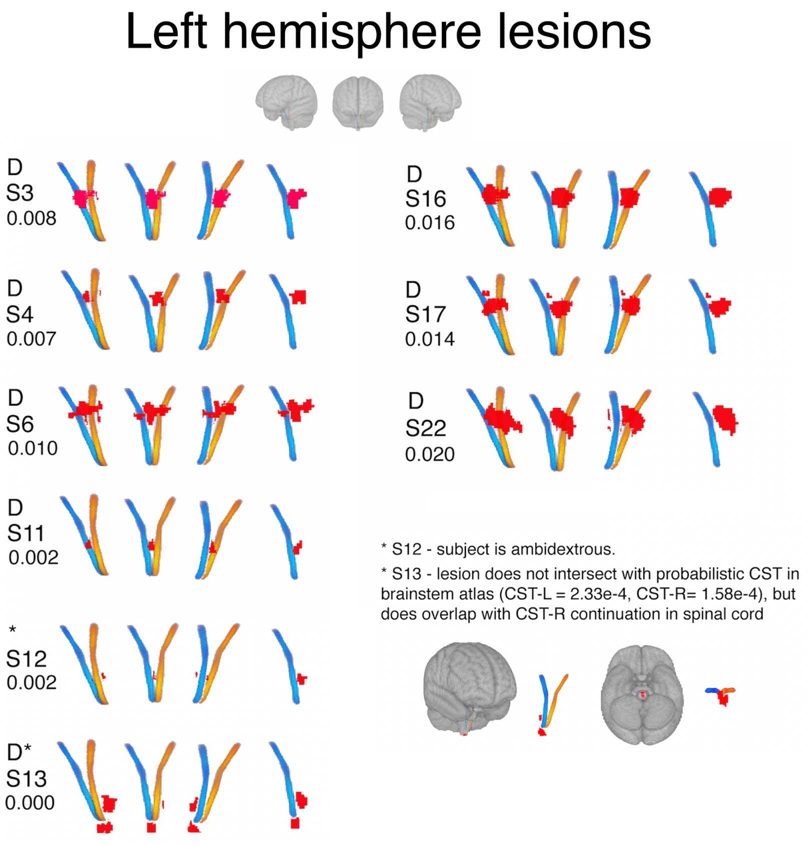


Figure S7: Left hemisphere lesion location relative to brainstem corticospinal tracts. Red = lesion. Blue = right CST, yellow = left CST. Four views of the lesions/CSTs are displayed, from left to right: left lateral view, anterior view, right lateral view. A "D" above the subject identifiers ("SX") indicates that the lesion is in that subject's dominant hemisphere. Numbers below subject identifiers indicate Dice overlap between lesion and ipsilesional CST.

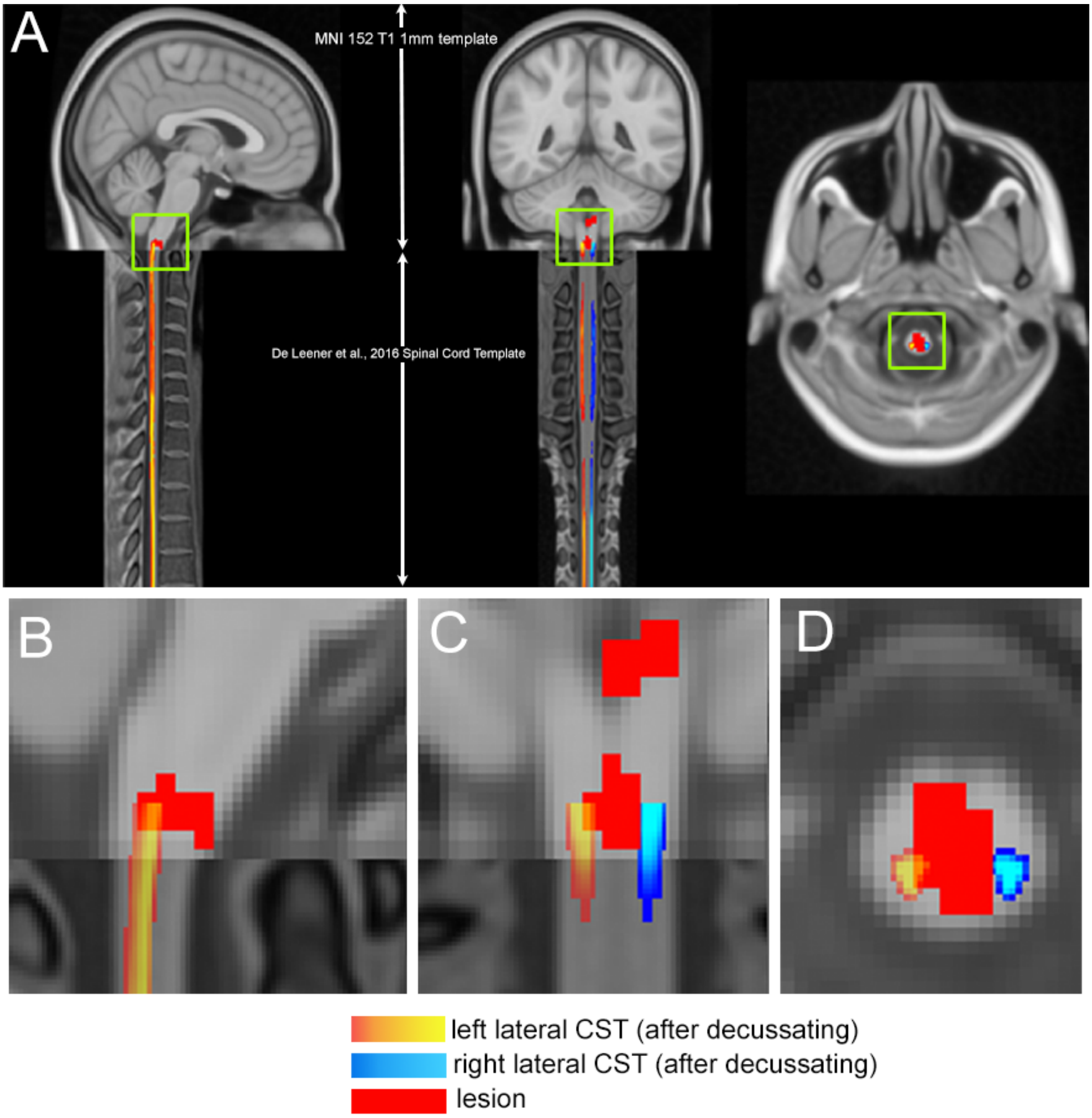


Figure S8: A.Sagittal, coronal, and horizontal views of the lateral corticospinal tract atlases and lesion for subject S13 (who is right-handed) on top of the MNI 152 brain template (top) and De Leneer et al. spinal cord template. Close-ups of B., sagittal slice, C., coronal slice, and D., horizontal slice with reduced opacity of the left lateral CST, which is the dominant CST as the subject is right-handed and the lesion is in the right spinal cord after decussating in the medulla.

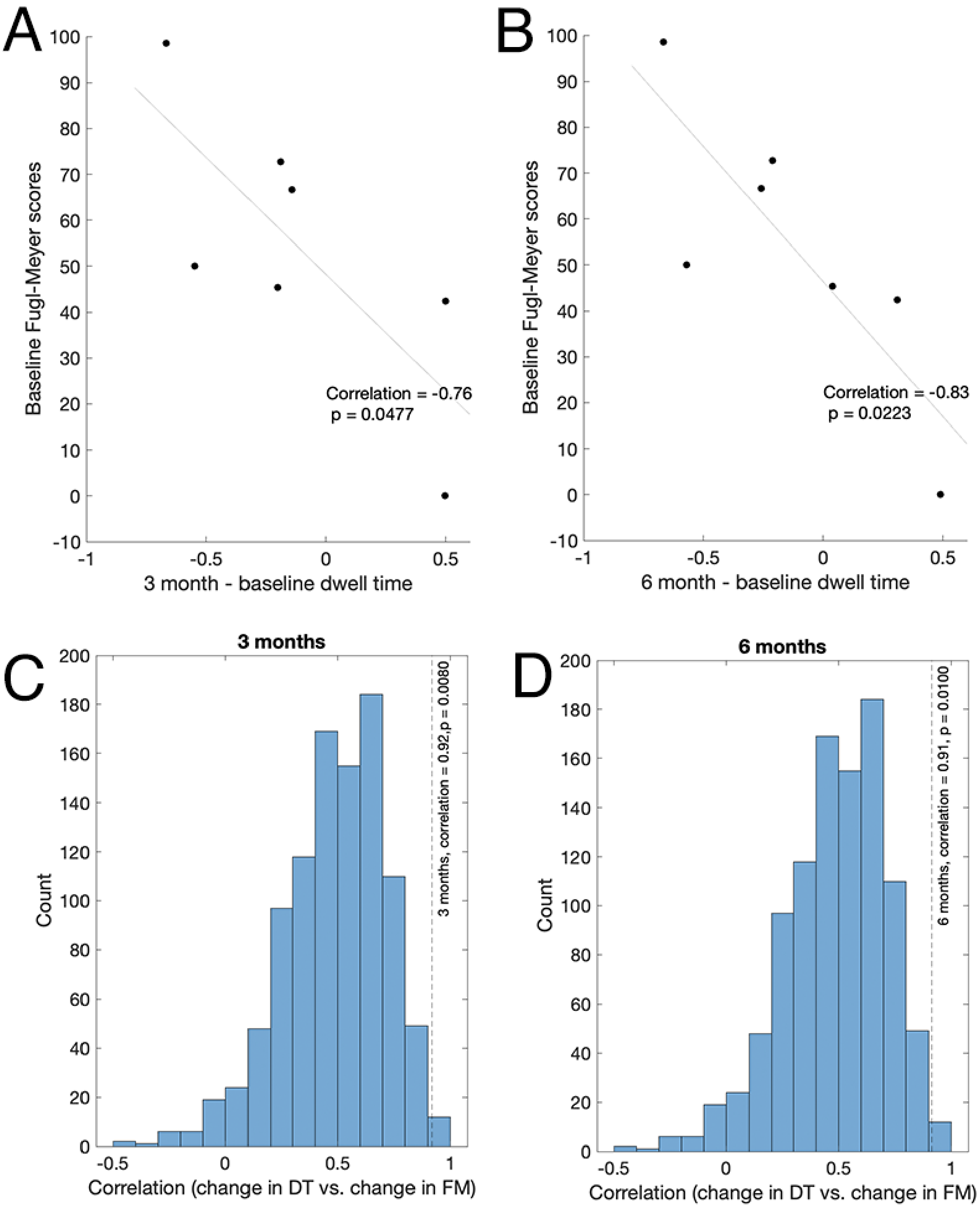


Figure S9: **A.** Correlation between 1-week (baseline) Fugl-Meyer scores, and the change in dwell time in $FPN^+$ between 1 week and 3 months in subjects with dominant hemisphere CST damage. **B.** Correlation between 1-week (baseline) Fugl-Meyer scores, and the change in dwell time in $FPN^+$ between 1 week and 6 months in subjects with dominant hemisphere CST damage. **C.** Distribution of correlations between change in dwell time and change in Fugl-Meyer using observed 1-week and 3-month dwell times, observed 1 week Fugl-Meyer scores, and 3-month Fugl-Meyer scores created according to the proportional recovery plus random noise (100 sets of 3-month Fugl-Meyer scores were generated). Observed correlation using observed 3 month Fugl-Meyer scores is shown as the grey dotted line (R = 0.92). **D.** Distribution of correlations between change in dwell time and change in Fugl-Meyer using observed 1-week and 6-month dwell times, observed 1 week Fugl-Meyer scores, and 6-month Fugl-Meyer scores created according to the proportional recovery plus random noise (100 sets of 6-month Fugl-Meyer scores were generated). Observed correlation using observed 6 month Fugl-Meyer scores is shown as the grey dotted line (R = 0.91).

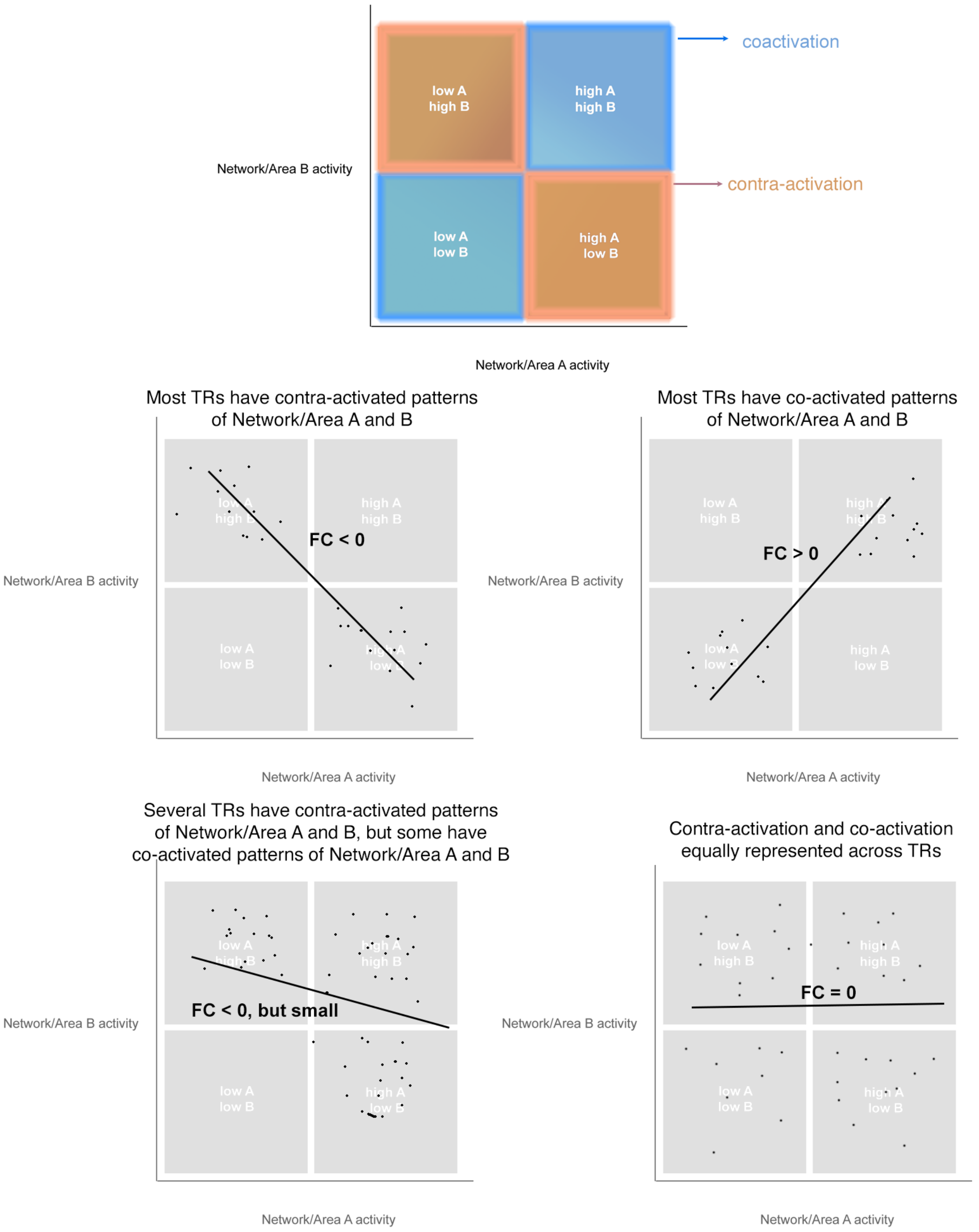


Figure S10: Theoretical framework for linking fractional occupancy of brain states to functional connectivity between brain regions. Each TR can be plotted into one of four quadrants representing the joint activity patterns of a given pair of networks: high activity of networks/regions A and B (coactivation); low activity of A and B (coactivation); high A, low B (contra-activation); and low A, high B (contra-activation). When most TRs are in a contra-activation configuration of 2 areas, then the functional connectivity between those areas will be negative; when most TRs are in a co-activation configuration, the FC between those two areas will be positive.
